# Mild membrane depolarization in neurons induces immediate early gene transcription and acutely subdues responses to successive stimulus

**DOI:** 10.1101/2020.07.23.217844

**Authors:** Kira D. A. Rienecker, Robert G. Poston, Joshua S. Segales, Ramendra N. Saha

**Author notes:** To whom correspondence should be addressed: Ramen Saha, Ph.D.: University of California, Merced, Room 346 S&E Building 1. 5200 North Lake Road, Merced, CA, USA, 95343.

## Abstract

The transcriptional profile of immediate early genes (IEGs) is indicative of the duration of neuronal activity, but it is unknown whether it affected by the strength of depolarization. Also unknown is whether an activity history of graded potential changes influences further neuronal activity. In this work with dissociated rat cortical neurons, we found mild depolarization – mediated by elevated extracellular KCl – not only induces a wide array of rapid IEGs, but also transiently depresses transcriptional and signaling responses to a successive stimulus. This latter effect was independent of *de novo* transcription, translation, calcineurin (CaN) signaling, and MAPK signaling downstream of PKC. Furthermore, as measured by multiple electrode arrays, mild depolarization acutely subdues subsequent spontaneous and bicuculline-evoked activity. Collectively, this work suggests that a recent history of graded potential changes acutely depresses neuronal intrinsic properties and subsequent responses. Such effects may have several potential downstream implications, including reducing signal-to-noise ratio during Hebbian plasticity processes.

---

In the brain, sensory stimulation and learning events alter neuronal activity and upregulate expression of immediate early and other genes, a phenomenon referred to as excitation-transcription coupling (1–3). Such activity-dependent transcription of immediate early genes (IEGs) is functionally important for adaptive processes such as memory consolidation, cognitive flexibility, Hebbian plasticity, and neuronal homeostasis (4–8). Different modes and patterns of stimulation induce distinct gene expression programs (9, 10). In fact, the transcriptional profile of the neuron is indicative of the duration of activity (11). We have shown activity-induced IEG expression is characterized by three waves, including rapid and delayed immediate early genes (rIEGs and dIEGs) as well as *de novo* translation-dependent Secondary Response Genes (SRGs). Sustained neuronal activity induces rIEG, dIEGs, and SRGs, whereas brief activity induces only rIEGs (11). These findings suggest that neurons can sense and respond to distinct activity patterns, such as its duration, with signature transcriptional programs.

While exact roles of IEGs remain unclear in learning and memory, several IEGs–such as *Arc, Npas4, c-Fos*, and *Egr1–* are often used to demarcate active neurons allocated to a memory trace or engram during Hebbian learning (1, 5, 12, 13). Interestingly, only a subset of neurons responds to sensory stimuli and are incorporated in engrams. Usually, these neurons are marked by enhanced excitability and IEG transcription. Because the size of an engram is limited, neurons are forced to compete for allocation (12, 14, 15). This competition is partially modulated by the activity of the transcription factor CREB, which regulates certain subsets of IEGs (16, 17) and bidirectionally modulates neuronal excitability (12, 18). Overexpressing CREB enhances a neuron’s competitiveness for allocation to an engram (14, 18–22), while neurons with decreased CREB function are more likely to be excluded (12, 14). Previous activity experience also impacts a neuron’s competitiveness. A successfully allocated neuron remains excitable for about six hours after a learning event, and during this period, it is likely to be co-allocated to a new engram representing a second event (18, 20). After longer periods, allocated neurons become ‘refractory’ or less excitable, and thereby less likely to be co-allocated to a second event (20). Together, it has been proposed that a neuron’s inclusion in engrams is in part determined by its intrinsic properties (19), which in turn depends on its recent history of activation (23).

The activity history of a neuron may include suprathreshold membrane polarization resulting in orthodromic or backpropagating action potentials, and also, subthreshold graded potential changes. While action potentials are traditionally viewed as key mediator of neuronal operations, studies have shown that subthreshold graded changes in membrane potential can effectively recruit neuronal second messengers (24), alter intrinsic properties (25), and modulate synaptic communication (26–28). Most activity-induced gene transcription studies have involved suprathreshold stimulation, including direct membrane depolarization using 55mM extracellular potassium chloride (KCl), homeostatic potentiation of synaptic activity by prolonged Tetrodotoxin (TTX) treatment followed by its washout, or disinhibition of inhibitory synapses using Bicuculline (Bic) (11, 29). Other approaches have used optogenetics to mimic stimulation parameters. With these tools, Yu *et al* determined one nuclear calcium transient induced by a single burst of action potentials is the minimum signal strength required to induce activity-dependent transcription in hippocampal neurons (30). However, it remains largely unknown if mild graded potential changes are capable of triggering transcriptional responses in neurons.

In the current study, we use rat dissociated cortical cells and low concentrations of extracellular KCl to first address whether neurons undertake IEG transcription in respond to mild depolarization. We also address whether such depolarization ‘experience’ leaves a cellular ‘memory’ to affect future transcriptional and electrical responses. Hereby, we present data to suggest that neurons presented with mild stimulation respond transcriptionally with distinct IEG profiles for various doses of KCl. When these mildly depolarized neurons are activated again after some time, they manifest subdued transcriptional, signaling, and electrical responses.

## Materials and Methods

### Tissue Culture

Primary neuronal culture in these experiments was performed as previously described (UC Merced IACUC approval: AUP#16-0004) (31). Samples were cortical neurons from E18 embryos plated on 35 cm2 dishes and cultured in Neurobasal feeding media with B27–Neurobasal medium (Gibco, #21103049), 25µM glutamate (Sigma-Aldrich, #1446600), and 0.125x B27 supplement (Gibco, A35828-01). Cells were maintained at 37°C in a humidified incubator with 5% CO2. Half the feeding media was replaced every 3-4 days. Neurons were used for assays between 10-14 days *in vitro*.

### Two-step paradigm

Isosmotic KCl stock solutions for 1° treatment were prepared at salt concentrations mimicking neurobasal media where increasing amount of KCl was compensated by decreasing NaCl. Stock solutions (pH 7.4) contained: variable KCl (5mM, 15mM, 35mM, 55mM; Fisher, P217), 1.8mM CaCl (Fisher, C79), 0.8mM MgCl (Fisher, BP214), 26mM NaHCO3 (Fisher, S233), variable NaCl (Fisher, BP358), 1mM NaH2PO4 (Fisher, BP330), 25mM D-Glucose (Sigma, G5767), and 11mM HEPES (Sigma, H3375).

Many experiments conducted for this paper were modified versions of our basic two-step protocol. In this protocol, a variable term 1° KCl treatment was followed by a variable term washout stage, and then a 15’ 2° treatment of 5µM bicuculine. To prepare the 1° KCl treatment, 500 µl of conditioned media was removed from the dish and pre-mixed with 500 µl of the appropriate KCl stock solution in artificial cerebrospinal fluid (ACSF) to achieve the final KCl concentration for application to the cells. KCl solutions were prepared before starting any treatments, and dishes were returned to the incubators while the remaining preparations were completed for treatment. When treatments were ready to begin, no more than 4 dishes were removed from the incubators at a time. Dishes were treated sequentially, and the researcher noted the exact time of treatment for each dish to keep the timing accurate. When treatment began, old media was removed from the dish with a micropipette and saved. Warm KCl treatment was added to the dish as soon as possible after old media was removed. Dishes were returned to the incubators for the time of KCl treatment. Dishes were removed from the incubator just prior to completion of the 1° stage, and KCl was removed on schedule using a micropipette. For this wash out stage, 1ml of old media from the original dish replaced the KCl solution, and 1ml of warm conditioned media was added to further dilute any residual KCl. Conditioned media consisted of media drawn on the same day from cultured primary cortical cells the same age and source as the treated cells. Dishes were returned to the incubator for the duration of the wash out stage. Dishes were removed from the incubator 2-3 minutes before the end of the wash out stage, and media was measured and reduced to 1ml per dish to prepare for 2° treatment. On schedule, 1µl DMSO or 5µM bicuculine was added to the 1ml media in the dish for the 2° treatment stage. Each dish was swirled to mix the solution after the addition of 2° treatment, and the collection of treated dishes was returned to the incubator for the duration of the 2° treatment time. Dishes were removed from the incubator just prior to the end of the 2° treatment time, media was removed via suction on completion, and appropriate sampling was performed.

### Treatments

Bicuculline and 4AP: Neurons were treated with 5µM Bicuculline (Bic; Sigma-Aldrich, #14340) to inhibit GABAergic activity in two-step treatments. The health of the cell culture was assessed in a test sample by treatment with 50µM Bic treatment and 75µM 4-Aminopyridine (4AP; Acros Organics, #104571000).

PMA and TTX: MAPK pathways were activated intracellularly via PKC with 1µM-1nM phorbol 12-myristate 13-acetate (PMA; Sigma-Aldrich, #P1585). PMA treatments were applied as a 2° treatment in place of bicuculine. PMA was diluted in dimethylsulfoxide (DMSO, Sigma #D2650) to achieve the variable PMA concentrations, so that the same DMSO load was added to each treated condition. PMA treatments were combined with 1µM TTX (Calbiochem, #554412) to block membrane activity.

CHX, DRB/CHX, and Flavopiridol (FP): Transcription and Translation inhibitors were applied from the beginning of treatments at time = 0 and were maintained at the same concentration through all stages of the experiment. Experimental designs contained both inhibitor-added and inhibitor-free conditions for each biological replicate. Flavopiridol aka FP (Sigma, #F3055), Cyclohexamide aka CHX (Sigma, #C7698), DRB (DRB; Sigma-Aldrich, #D1916).

FK506: FK506 aka FK (Tocris, #3631) treatments (1µM) were applied from time = 0 at the primary stage for “FK primary” condition, and from time = 1 hr 30 min at the start of the secondary stage for the “FK secondary” condition. A control condition was run alongside these treatments, and DMSO was added at each stage of treatment to keep total DMSO additions equivalent across all three conditions. Therefore, all three conditions contained 1µl DMSO per ml from time = 0 through 1 hr 30 min at the start of the 2° stage, and 3µl DMSO per ml from there on, as the FK secondary condition received FK in 1µl DMSO and all conditions received 1µl 5µM Bic or DMSO as the 2° treatment.

### Electrophoresis and Western blotting

Samples were lifted from cell culture dishes with 75 µl 1x RIPA, made in-house (25mM Tris-HCl pH 8; 150mM NaCl; 1% Na-deoxycholate; 0.1% SDS; 0.1% IGEPAL) supplemented with 1:100 protease/phosphatase inhibitor cocktail (Thermo, #78442). Lysates were sheared by sonication(3 x 30 s, lowest setting on Bioruptor®). Cell debris were pelleted at 15,000 rpm for 5 minutes at 4°C, and clarified supernatant was removed to a new 1.5ml tube. For western preparation, equal volumes of the supernatant were used for each sample. Samples were combined with Dye (4x Laemeli buffer (BIO-RAD, #1610747) with 10% betamercaptoethanol (Sigma, 63689) and were boiled for 5 minutes at 95° in a heat block. Sample and dye mixtures were then loaded in a 4 – 20% (BIO-RAD, #4568095) or 4-15% (BIO-RAD #456-1083) Mini PROTEAN® gel in Tris/Glycine/SDS (BIO-RAD #1610772). Gels were run at 150 V for ∼10 minutes and then 110 V until the dye band reached a few centimeters above the end of the gel. Resolved proteins were transferred to a PVDF membrane (BioRad, 10026933) using the BioRad Trans-Blot Turbo Transfer System with 20% EtOH-containing trans-blot turbo transfer buffer (BioRad, #10026938) on the mixed molecular weight setting (7 minutes). PVDF membranes were immediately transferred to cold TBST, and were incubated at 4°C overnight in 1° antibody in 5ml 1x TBS-T with 1.5% BSA (Fisher, BP9703). The pERK antibody (Rabbit, Cell Signaling, 4370S) was diluted at 1:1000, and the bet-actin antibody (Mouse, Thermo Fisher, AM4302) was diluted at 1:10,000. Membranes were washed three times in 1x TBS-T before being probed with 2° antibody for 45 minutes at room temperature. 2° antibodies were either goat-anti-Mouse 647 (RRID: AB 2535808) or goat-anti-Rabbit 546 (RRID: AB 2534093) Alexa Fluor secondary antibodies (Life Technologies). Membranes were washed three times with 1x TBS-T for 5 min each, and imaged using BIO-RAD Multiplex ChemiDocTM Imaging System.

### RNA extraction and gene transcription quantitation with real-time PCR

Total RNA was collected using the illustra RNAspin Mini kit (GE Lifesciences, #25050072). Samples were collected with 350 µl RNA lysis buffer from the kit, and after processing were precipitated in 40 µl RNase-free water. Specific pre-mRNAs from total RNA samples were initially amplified by cDNA synthesis (14 cycles) using primers overlapping an intron-exon junction and a OneStep RT-PCR kit (Qiagen, #210212). Each reaction used 250 ng of RNA per reaction. The housekeeping transcript was Gapdh. The cDNA product was diluted 1:20 with RNase-free water and 4 µl were used for each quantitative real-time PCR (qRT-PCR) using PerfeCTa SyBR Green FastMix (QuantaBio, #95072-012) and the BIO-RAD CFX Connect realtime PCR Detection System. Samples were run in technical duplicates for each primer, and the average Ct value was used with the ΔCt method to calculate fold-change.

### Hierarchical clustering of rapid immediate early gene expression

Pre-mRNA levels of rapid immediate early genes under various experimental conditions were used for hierarchical clustering analysis to summarize results and reveal trends in regulation of gene expression. Each gene was represented as a vector of fold-change values (for each time point and treatment group) and organized into matrices for clustering analysis. Specific normalization details are provided in figure legends. Matrices were uploaded to Morpheus (https://software.broadinstitue.org/morpheus), a tool made available through the Broad Institute, which was used to generate expression heatmaps, perform clustering analysis, and export dendrograms. Briefly, the clustering analysis computes the Euclidian distance between genes based on provided features (expression levels under various conditions and time points) which are then used to recursively pair genes into clusters by average linkage, from closest to farthest, generating dendrograms to visualize relatedness.

### Microelectrode array (MEA) experiments

Neurons from the preparations described in the cell culture methods section were plated on poly-L-lysine/laminin coated MEAs (60MEA200/30-Ti, Multi Channel Systems (MCS), Reutlingen, Germany) in 600μL of B27-supplemented Neurobasal plating media. Cells were fed every 3-4 days by exchanging approximately half the media with B27-supplemented BrainPhys feeding media (StemCell). This was done to promote optimal conditions for neuronal firing which has been shown to be enhanced in BrainPhys media (32). Recordings were made with an MEA2100-lite system that interfaces with MCS provided Multi Channel Experimenter software. Sampling was conducted at 10-kHz in 2-minute sessions at room temperature (arrays were covered to prevent contamination).

### Microelectrode array (MEA) data analysis

Recording were initially post-processed in Multi-Channel Analyzer with a high-pass 1st order Butterworth filter with 100hz cutoff prior to generation of spike time-stamps. Spikes were detected using an automatic threshold estimator set to 5-8 standard deviations from the baseline signal depending on the amount of baseline noise. To quantify burst properties, the Multi-Channel Analyzer burst detection tool was used with the following settings: max. interval to start burst, 25ms; max. interval to end burst, 250ms; min. interval between bursts, 500ms; min duration of burst, 50ms; min. number of spikes in burst, 5. Raster images from example recordings were generated in NeuroExplorer (Nex Technologies, Colorado Spring, Colorado USA). For data presented in Figure 6, each recording was summarized as an average of all electrodes for various parameters (Number of spikes, etc.). For data in Figures 7 and 8, electrodes were individually analyzed and displayed, pooling all electrodes for each treatment group across replicates to generate the distributions presented. Individual electrode data was generated from Multi Channel Analyzer and then further processed in R (R Core Team, 2014). Plots and statistics were generated using GraphPad Prism version 8.4.2 (GraphPad software, San Diego, California USA).

### Statistics of qRT PCR and Westerns

Data were analyzed using GraphPad Prism 7 (RRID: SCR_002798, GraphPad software, San Diego, CA). Where possible, normality was assessed with the Shapiro-Wilk test and was reported, though ANOVA was carried out regardless as it is somewhat robust to deviations from normality. Outliers were identified for each cell of the design using ROUT, Q = 5%. All analyses were run with and without outliers. Results for all outliers included are reported, and any differences when outliers were excluded are noted. Data were analyzed with appropriate 3-way, 2-way, or 1-way ANOVAs. Three- and two-way interactions were considered first. When insignificant, we next considered two-way interactions and main effects respectively. When interactions were significant, we re-organized the data to investigate simple main effects by collapsing over insignificant factors and/or by separating the data by each level of one factor for analysis. Prism does not automatically use the pooled error term from the larger ANOVA comparison, so for main effects and simple main effects analyses following up on larger ANOVAs, we ran the reorganized data using the next ANOVA down. For example: for 3-way ANOVAs with insignificant 3-way interactions but a significant 2-way interaction, we reorganized the data by collapsing over the insignificant factor and ran an explicit 2-way ANOVA. We then followed the same procedure for the 2-way ANOVA, running a one-way ANOVA or unpaired t-test for simple main effects at each level of a factor if the 2-way interaction was significant, or, for main effects, by collapsing the data again over the insignificant factor and running a one-way ANOVA or unpaired t-test on the entire data set. Multiple comparisons were reported for the lowest-level ANOVA and the Bonferroni correction (CHECK) was applied to adjust the significance. Mean differences are reported with the 95% confidence interval and significance value. Alpha levels were set to 0.05. Error bars represent standard error of the mean throughout, except where otherwise noted. Biological replicates are indicated throughout as N in corresponding figure legends. Biological replicates constitute cell culture preparations from the pooled cortices of embryos from independent litters.

## Results

### Variable KCl induces IEG pre-mRNA expression

We first treated primary cortical neurons with variable mild KCl treatments (below field standard of 50-55mM) for thirty minutes. While a few publications have used mild KCl treatments (33– 39), these publications have focused on specific IEGs, such as *Bdnf* (37) and *cFos* (39), and only a handful have used treatment times under an hour (33, 35, 37). Here, we surveyed 15 rapidly induced IEGs (11) for multiple timepoints within the hour to study induction and its dynamics. Fourteen of fifteen genes were significantly induced by 50µM bicuculline and 75µM 4AP treatment (Bic+4AP). 30mM KCl induced eight rIEGs, 20mM KCl induced eleven rIEGs, and 10 mM KCl induced two rIEGs (Fig 1A). In all cases, induction under treatment with 5mM KCl (same KCl concentration as the commercial media) was not significantly different to mechanical controls. We then used clustering analysis to determine which IEGs responded most similarly across variable KCl treatments (Fig 1B). While some IEGs were similarly induced, many showed different responses to 20mM and 30mM KCl. For instance, 20 mM KCl significantly induced *Npas4, Cyr61, Dusp1*, and *Fbxo33* above mechanical controls (M), while 30 mM KCl did not. We concluded mild treatments of 20-30mM KCl for 30 minutes are sufficient to significantly induce most IEGs, but 20 and 30mM KCl may have overall different induction profiles.

Next, we selected *Arc* as an example to investigate the effect of mild KCl treatments on IEG induction over time. Compared to 5mM KCl treatment, as reported before, *Arc* was significantly induced by 20mM KCl – similar to Bic+4AP (40)– at 15’, 30’, 45’, and 60’, with a peak at 30’. Although 30mM KCl induced *Arc* at 30’ and 60’, the response profile was much flatter (Fig 1C). Noticeably, increasing the concentration of KCl did not necessarily induce more transcription, or more closely follow the Bic+4AP induction profile. By visual inspection, 20mM KCl had the most similar induction profile to Bic/4AP for *Arc*. These time course data indicate that mild KCl treatments can induce IEG transcription, and suggests neurons respond to different concentrations of KCl with different transcriptional profiles over time.

We also tested whether an extremely short KCl treatment induces *Arc*. We treated primary cortical cultures with 1’ of variable KCl, followed by a wash out step and sampling at 10 or 15 minutes to allow transcription mechanisms time to react. Wash out involved replacing KCl treatment with conditioned media saved from the same dish prior to KCl treatment. 10 mM, 20 mM, and 30 mM KCl all succeeded in inducing significantly more *Arc* pre-mRNA than the mechanical control (Fig 1D). We concluded, even 1’ of mild KCl can elevate *Arc* transcription 10 to 15 minutes later.

**Figure 1.**
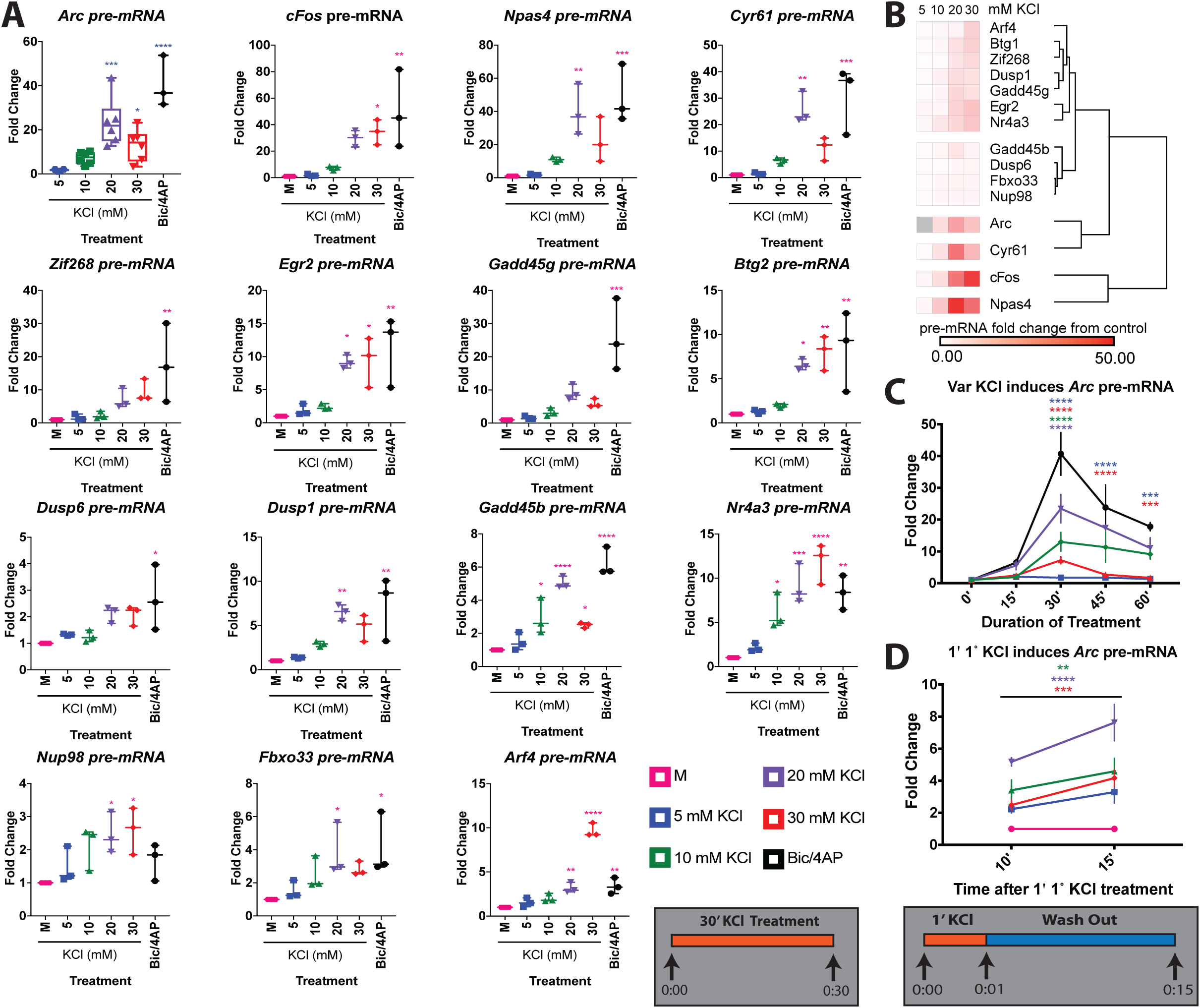
30’ Treatment with Variable KCl induces IEGs. **(A-B)** Samples were treated with variable KCl for 30 minutes and IEG pre-mRNA was quantified. N= 3 for all conditions for all IEGs except *Arc* pre-mRNA, for which there was N = 6 for 5, 10, 20, and 30 mM KCl, and N = 3 for the Bic/4AP treatment. Details of one-way ANOVAs are found in Table I. **(A)** All IEGs showed induction by Bic/4AP except *Nup98*. 30 mM KCl induced 8 IEGs, 20 mM KCl induced 11 IEGs, and 10 mM KCl induced 2 of the 15 IEGs tested. 5 mM treatments were not significantly different to mechanical controls. **(B)** IEGs were clustered by the similarity of their induction over all variable KCl treatments. **(C)** Samples were treated with variable KCl for variable time and IEG pre-mRNA was quantified. N = 3 for 0 ‘, 45’, and 30’ Bic/4AP; N = 4 for 60’, 15’ 5mM; N = 5 for 15’; N = 6 for 30’. For *Arc* pre-mRNA: Interaction (F(16,76) = 5.842, *P* < 0.0001); TIME (F(4,76) = 34.97, *P* < 0.0001); TREATMENT (F(4,76) = 33.19, *P* < 0.0001). One-way ANOVAs were performed for the simple main effect of TREATMENT at each level of TIME, with comparisons made to the 5mM KCl treatment. At 15’, (F(4,19) = 6.595, *P* = 0.0017), Bic/4AP (*P* = 0.0067) and 20 mM KCl (*P* = 0.0325) induced more *Arc* pre-mRNA than 5mM KCl. At 30’ (F(4,22) = 18.04, *P* < 0.0001), Bic/4AP (*P* < 0.0001), 20mM (*P* = 0.0001), and 30mM KCl (*P* = 0.0469) induced significantly more *Arc* pre-mRNA than 5mM. At 45’ (F(4,10) = 6.055, *P* = 0.0097), only Bic/4AP (*P* = 0.0077) induced significantly more than 5mM. At 60’ (F(4,15) = 16.30, *P* < 0.0001), Bic/4AP (*P* < 0.0001), 20mM (*P* = 0.0038), and 30mM (*P* = 0.0191) induced more than 5mM. **(D)** Samples were treated with 1 minute of variable KCl and IEG pre-mRNA was quantified after 10 and 15 minutes. N = 3 for 10’ samples and N = 4 for 15’ samples. For *Arc* pre-mRNA: Interaction (F(4,25) = 0.9513, *P* = 0.4511); TIME (F(1,25) = 9.632, *P* = 0.0047); TREATMENT (F(4,25) = 18.34, *P* < 0.0001). Main effects analysis of TREATMENT showed 10mM (*P* = 0.0009), 20mM (*P* < 0.0001), and 30mM KCl (*P* = 0.0115) induced significantly more *Arc* pre-mRNA than the mechanical control. * indicates *P*-value < 0.05, ** < 0.01. *** < 0.001

### Rapid IEG transcription and MAPK/ERK pathway are depressed after 1° KCl treatment in a two-step paradigm

We designed a two-step treatment paradigm to investigate the effect of mild KCl treatment on neuronal responses to subsequent stimulation (Supplemental Fig. 1). Primary cortical cultures aged DIV10-14 were treated first with variable concentration of KCl for 30’ (1° KCl), which was then removed and replaced with conditioned media in a 1-hour recovery step (‘wash out’ in schema), followed by a 15’ 2° treatment with 5µM bicuculline (2° Bic) (Fig 2). We used 5µM Bic instead of 50µM Bic to avoid ceiling or floor effects on IEG transcription and compared 30mM KCl pre-treatment to mechanically handled (M) and 5mM KCl controls. We chose to use 30 mM KCl because it induced the highest *Arc* pre-mRNA levels at 30’ of treatment. *Arc* is used as a key indicator of rapid IEG induction for several subsequent experiments in this paper. Of the 15 IEG pre-mRNAs tested, 10 showed significant depression of IEG pre-mRNA after 1° treatment with 30mM KCl. For all genes except *Gadd45b, Nup98*, and *Arf4*, KCl doses (M, 5mM, and 30mM) had different effects at each level of BIC (Bic or DMSO). Simple main effects analyses for 2° Bic and DMSO treated samples are displayed in Table II. Main effects analyses for *Gadd45b, Arf4*, and *Nup98* are shown in Table III. These results showed for most, but not all, of the rapid IEGs tested, prior treatment with 30mM KCl depressed subsequent pre-mRNA induction, even after a 1-hour recovery period.

**Figure 2.**
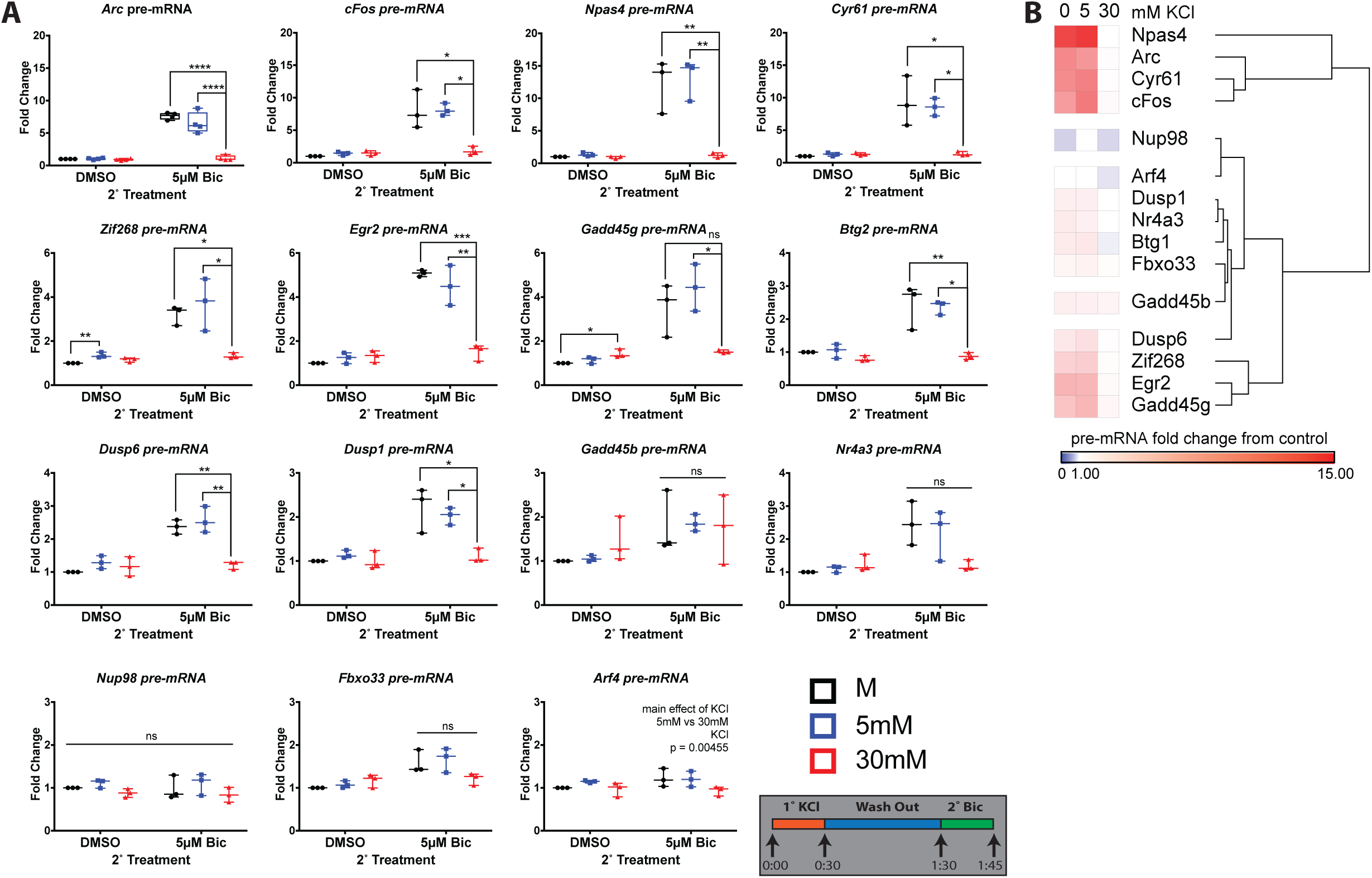
Rapid IEGs are depressed by 1° treatment with 30mM KCl. Rapid IEG induction was measured after the two-step paradigm. Most, though not all, rapid IEGs showed a depression in their transcription after pre-treatment with 30’ of 1° 30mM KCl, compared to samples treated with 5mM or mechanical controls. There was a significant interaction between BIC condition and KCL treatment for all genes except *Gadd45b, Nup98*, and *Arf4*. Significant interactions were followed up with simple main effects analysis via one-way ANOVA for KCL at each level of BIC. Most genes showed significant differences between 30mM KCl treatment and M or 5mM KCl for Bic treated samples but not for DMSO treated samples (exceptions *Gadd45g* and *Zif268*). Statistics details in results and Tables II and III. N = 4 for *Arc* and N = 3 for all other genes. * indicates *P*-value < 0.05, ** < 0.01. *** < 0.001, ns = not significant.

We then asked whether the duration of 1° treatment varied the depressive effect of 30mM KCl pre-treatment. For these and subsequent analyses, we focused on *Arc* as a representative IEG. Primary neuronal cultures were treated with variable 1° KCl for 1’, 15’ or 30’ intervals, before a one-hour recovery step and a 15’ 2° Bic treatment. Because we expect Bic but not DMSO to induce *Arc* pre-mRNA, we broke up the data by TREATMENT and use a two-way ANOVA to analyze the effects of KCL and TIME. For Bic treated samples there was a significant main effect of KCL and of TIME but no interaction. Both 30mM and 20mM 1° KCl treatment showed significant depression of transcription compared to controls (Figure 3A). We also found 30’ 1° treatment induced significantly less *Arc* pre-mRNA than 1’ 1° treatment or 15’ 1° treatment (Figure 3B). In contrast to Bic treated samples, DMSO treated samples had neither a significant interaction between KCL and TIME, nor any significant main effects of KCL or TIME. We concluded the depressive effect of 30mM 1° KCl treatment on subsequent induction of IEG pre-mRNA persisted despite time of treatment (down to 1’ treatment), but samples with 30’ pre-treatments induced overall less pre-mRNA than 1’ or 15’ conditions.

**Figure 3.**
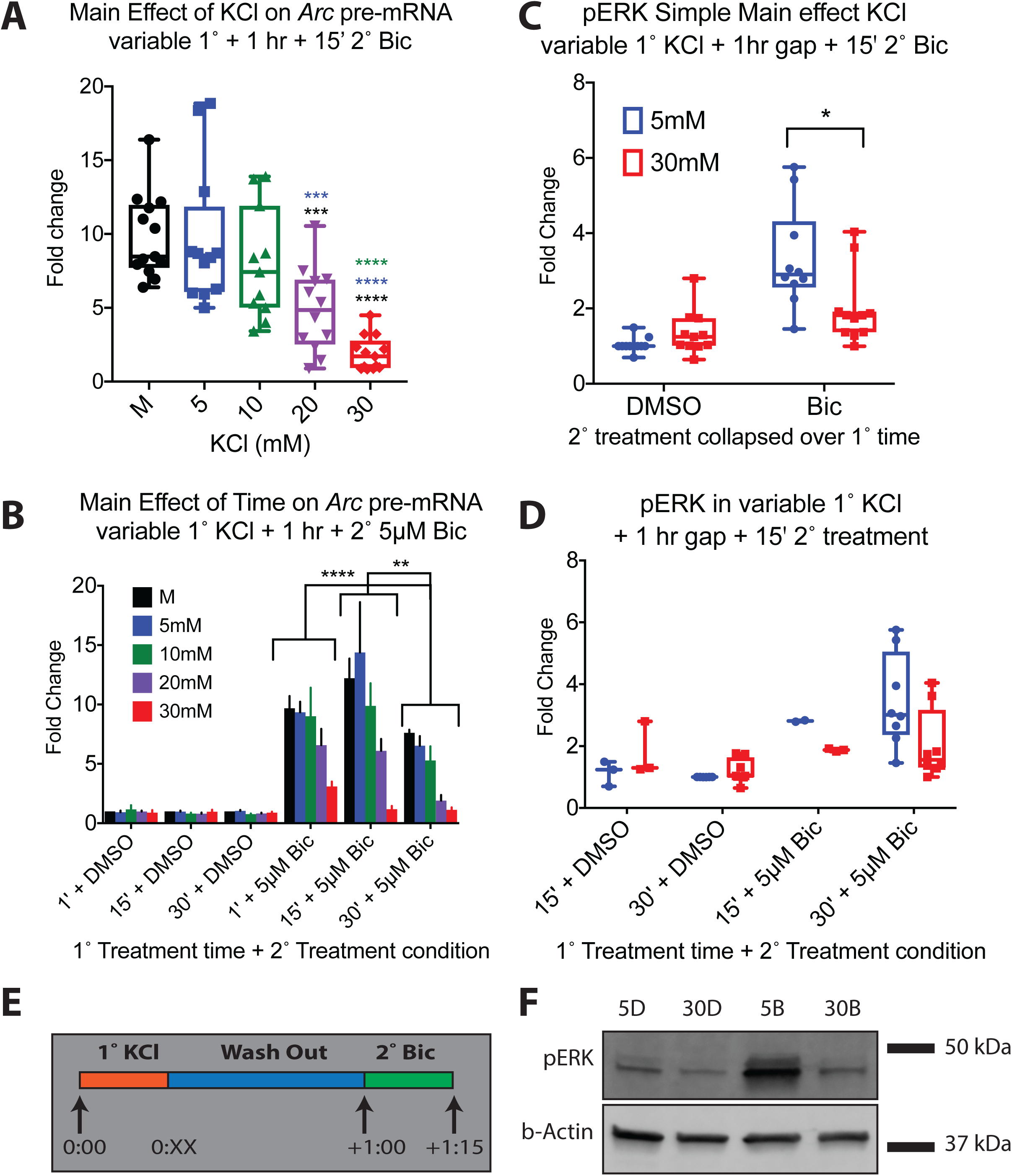
*Arc* pre-mRNA under Variable 1° KCl + 1 h + 2° 5µM Bic. Primary neuronal culture was treated with KCl for variable 1° treatment duration before 1-hour washout and a 15’ 2° 5µM Bic treatment. **(A-B)** qPCR data were analyzed with two-way ANOVA. For Bic treated samples: Interaction (F(8,46) = 1.464, *P* = 0.1967); KCL (F(4,46) = 22.34, *P* < 0.0001), TIME (F(2, 46) = 13.50, *P* < 0.0001). N = 3 for 1’ 1° 10mM KCl; N = 5 for 1’ 1° 20mM KCl; N = 4 for all other KCl conditions and treatment times. **(A)** The main effect of KCl for Bic treated samples, collapsed over TIME. Samples treated with 30mM KCl in the 1° stage induced significantly less *Arc* pre-mRNA than mechanical (*P* < 0.0001), 5mM KCl (*P* < 0.0001), and 10mM KCl (*P* < 0.0001). 20mM 1° KCl treatment also depressed *Arc* pre-mRNA compared to mechanical (*P* = 0.0002) and 5mM KCl (*P* = 0.0005). **(B)** The main effect of TIME collapsed over KCL. Bic treated samples treated for 30’ in the 1° stage induced significantly less *Arc* pre-mRNA over all levels of KCL than samples in treated for 1’ (*P* = 0.0021) or 15’ (*P* < 0.0001). There was no significant interaction between KCL and TIME for samples treated with DMSO during the 2° stage, nor were there any significant main effects. **(C-D)** 3 way interaction (F(1,35) = 0.0003, *P* = 0.9867); BIC x TIME (F(1,35) = 1.442, *P* = 0.2379); KCL x TIME (F(1,35) = 0.5238, *P* = 0.4740); BIC x KCL (F(1,35) = 6.323, *P* = 0.0167). N =3 for 15’ 1° treated samples; N = 8 for 30’ 1° treated samples. **(C)** Simple main effect of KCL collapsed over time (Interaction (F(1,39) = 9.553, *P* = 0.0037)). For Bic treated data there was a significant reduction in pERK levels after 30mM KCl treatment compared to 5mM KCl (*P* = 0.0171). There were no significant differences in pERK level between DMSO treated samples. **(D)** Total data for pERK by variable time. There was no significant effect of time. **(E)** Treatment diagram for variable 1° treatment. **(F)** Western blot of pERK induction for 30’ 1° treatment. DMSO treated samples induced little pERK. For 2° Bic treated samples, 1° treatment with 30 mM KCl reduced pERK induction compared to 1° treatment with 5mM. * indicates *P*-value < 0.05, ** < 0.01. *** < 0.001

Previously, we have shown that transcription of neuronal rIEGs relies on the MAPK/ERK pathway (11). Therefore, next, we investigated the impact of KCl pre-treatment on the magnitude MAPK/ERK pathway induction by assessing pERK levels. For these experiments, we used only 15’ and 30’ 1° KCl treatments (Fig 3D). We performed a three-way ANOVA on the variables TREATMENT_TIME (15’ 1° or 30’ 1°), BIC (DMSO 2° or Bic 2°), and KCL (5mM or 30mM). There was no significant three-way interaction, nor were there significant two-way interactions between BIC x TIME or KCL x TIME. The two-way interaction between KCL x BIC remained significant when we consolidated the data across TIME. For Bic treated samples, a t-test showed 30mM 1° KCl significantly depressed pERK compared to 5mM 1° KCl. There was no significant difference among DMSO treated samples. We concluded 30mM KCl pre-treatment significantly depresses both *Arc* pre-mRNA and pERK induction compared to controls, across all tested 1° treatment durations.

Next, we varied the time between 1° and 2° treatments to discover how long the effect of 1° KCl lasted. Primary neuronal culture was treated with KCl for a 1’ 1° treatment before a variable washout duration and a 2° 15’ 5µM Bic treatment. As before, we used a two-way ANOVA to analyze the effects of KCL (M, 5mM, 30mM) and TIME (1 hr, 2 hr, 4 hr) at each level of TREATMENT (Bic/DMSO) separately. For Bic treated samples, there was a significant two-way interaction between KCL and TIME. As before, 30mM 1° KCl depressed *Arc* pre-mRNA compared to 5mM and mechanical conditions for treatments with a 1 hour recovery period. In contrast, there was no significant main effect of KCl for samples with 2 hr or 4 hr wash out times (Fig 4A). 30mM KCl induced significantly less *Arc* pre-mRNA than mechanical and 5mM KCl controls, and this effect persisted even when outliers were removed.

**Figure 4.**
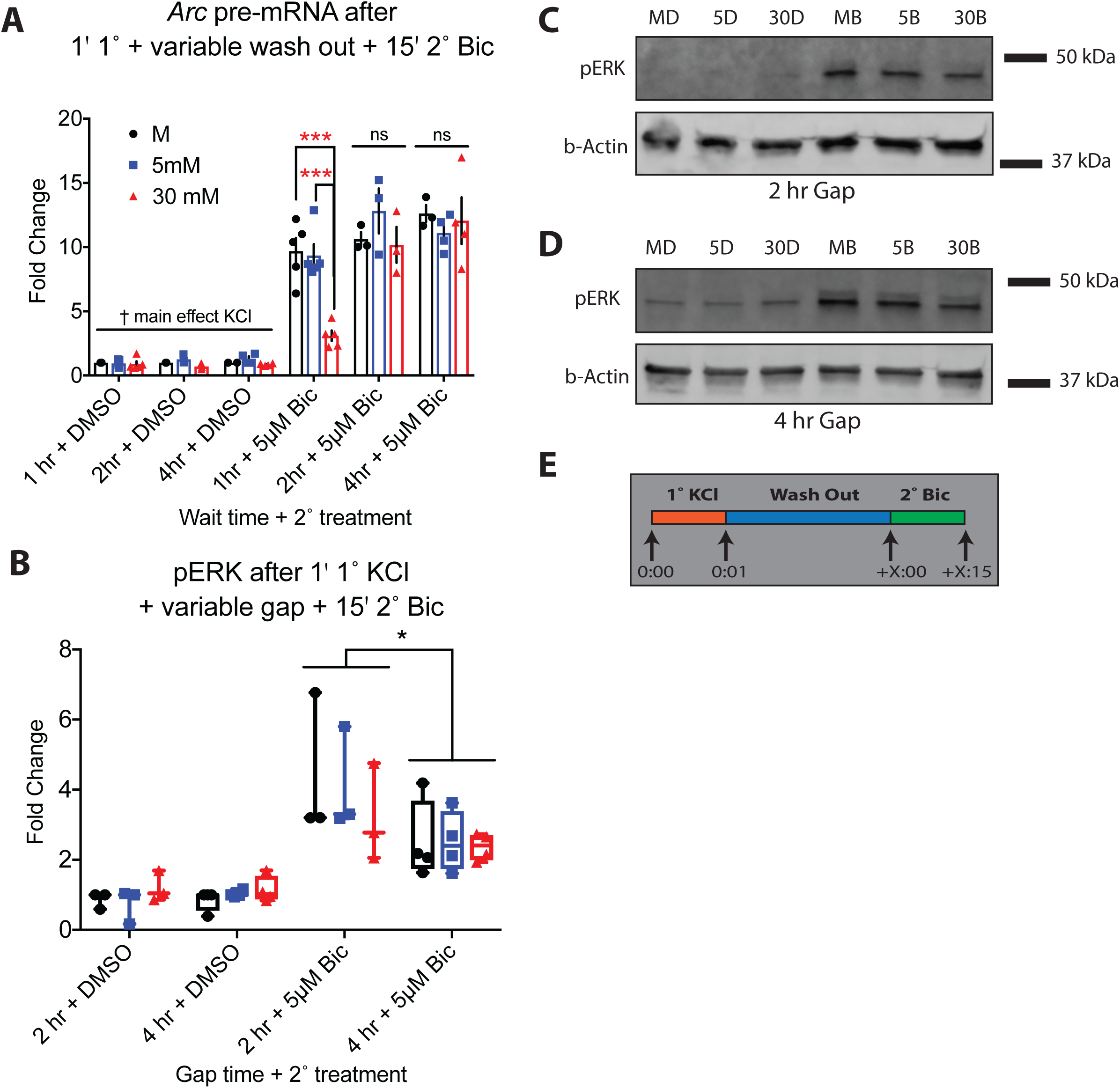
*Arc* pre-mRNA after 1’ 1° + variable wash out + 15’ 2° Bic. Wash out time between a 1’ 1° KCl treatment and a 15’ 2° 5µM Bic treatment was varied. **(A)** *Arc* pre-mRNA for Bic treated samples: Interaction (F(4,26) = 4.470, *P* = 0.0070); Simple main effect of KCL at 1 hr (F(2,12) = 20.67, *P* = 0.0001); at 2 hr (F(2,6) = 1.130, *P* = 0.3834); at 4 hr wait (F(2,8) = 0.3469, *P* = 0.7170). At a 1 hour wait, 30mM 1° KCl treated samples induced significantly less *Arc* pre-mRNA than both mechanical (*P* = 0.0003) and 5mM KCl 1° treated samples (*P* = 0.0004). After 2 and 4 hours, this depression of *Arc* pre-mRNA had disappeared. (For DMSO treated samples; Interaction F(4,25) = 1.151, *P* = 0.3561; TIME F(2,25) = 0.4571, *P* = 0.6383; KCL (F(2,25) = 4.558, *P* = 0.0205). 5mM and 30mM KCl treated samples were significantly different (*P* = 0.0154).) N = 5 for M, 5, and 30mM + 1hr; N = 4 for 5mM + 4hr and 30mM + 4 hr; N = 3 for M + 4 hr, M + 2 hr, 5mM + 2hr, and 30mM + 2hr. N = 2 for M + 4 hr DMSO sample. **(B)** pERK data was analyzed: 3-way interaction (F(2,30) = 0.4087, *P* = 0.6682); KCL x TIME (F(2,30) = 0.2623, *P* = 0.7711); KCL x BIC F(2,30) = 1.135, *P* = 0.3349; BIC x TIME F(1,30) = 6.945, *P* = 0.0132). When data were collapsed over KCL there was a significant two-way interaction between BIC x TIME (F(1,38) = 7.887, *P* = 0.0078). For Bic treated samples, pERK levels induced after a 2 hour gap were higher than those induced after a 4 hour gap (*P* = 0.0118). There were no significant differences between DMSO treated samples treated for 2hr or 4 hr. N = 4 for 4 hr samples; N = 3 for 2 hr samples. **(C)** Western blot of pERK induced after a 2 hr wash out step. **(D)** Western blot of pERK induced after a 4 hr washout step. There were no statistically significant differences between KCl conditions for either wash out time. **(E)** Treatment diagram for variable wash out time. * indicates *P*-value < 0.05, ** < 0.01. *** < 0.001, ns = not significant.

pERK data was analyzed with a three-way ANOVA to investigate the effects of KCL (5mM or 30mM), BIC (DMSO or Bic), and TIME (2 hr or 4 hr). Overall, the two-way interaction between BIC x TIME was statistically significant and remained so when we collapsed the data over KCL, for which there was no significant effect. pERK levels in Bic treated samples after a 2-hour gap was higher than after a 4 hr gap (Fig 4B). This difference did not exist in DMSO treated samples. We concluded the depressive effect of 30mM 1° KCl treatment on both *Arc* pre-mRNA and pERK was present after 1 hour recovery period, but disappeared by 2 hours. Notably, while equivalent levels of pre-mRNA were induced after 2 or 4 hours wash out, pERK induction was significantly lower after 4 hours wash out. Homeostatic recruitment of signaling cascades other than MAPK/ERK (like calcineurin signaling) may explain this difference between pERK and transcription activity.

### Depressive effects of 1° KCl treatment do not recover after de novo transcription or translation inhibition

To understand the underlying mechanism(s) of depressive effects produced by 30mM 1° KCl treatment, we next tested whether *de novo* translation and/or transcription are necessary for the effect. It is possible newly translated protein is responsible for the depressive outcome during the 2° induction. New protein might be translated from *de novo* transcripts induced by the 1° treatment, or from mRNA locally stored and translated at the synapse (41–43). This might result in short term negative feedback loops, which are known to exist in IEG regulation (e.g. *Sik1*) (44, 45). Therefore, we treated samples using our two-step paradigm with and without the translation inhibitor Cyclohexamide (CHX). *Arc* pre-mRNA data were first analyzed with three-way ANOVA and the factors KCL (mechanical handling aka M, 5mM, 10mM, 20mM, and 30mM 1° KCl), BIC (2° Bic or DMSO), and CHX [CHX(-) or CHX(+)]. Data are displayed in Figure 5A. The three-way interaction was not significant, but all three two-way interactions were significant. Therefore, we collapsed the data across single factors to analyze the two-way ANOVAs for simple main effects.

**Figure 5.**
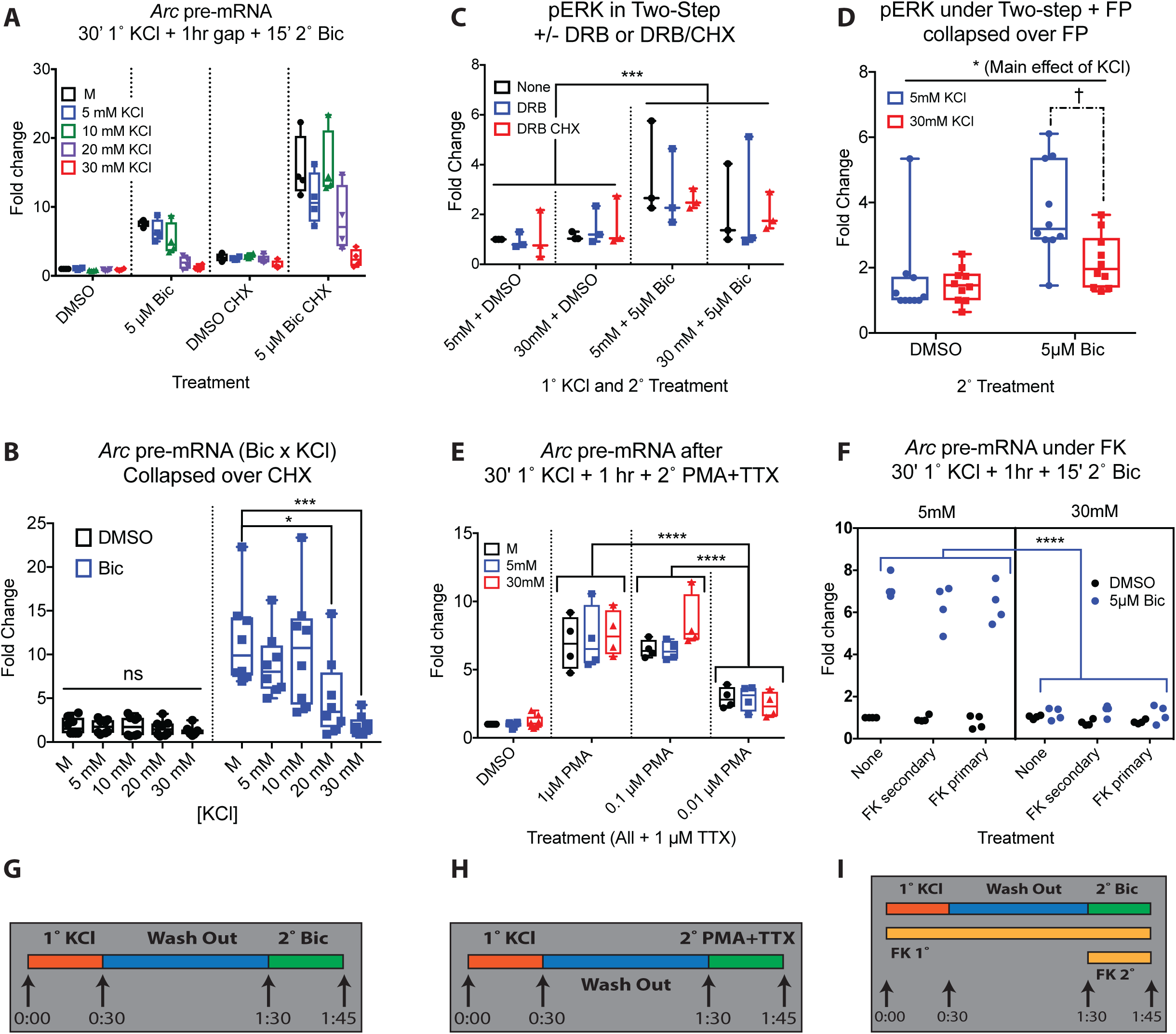
*Arc* pre-mRNA remains depressed under inhibition of translation. **(A-B)** Samples were treated with variable 1° KCl and 2° Bic with and without the translation inhibitor CHX. N = 4 for all cells of the design. All points are displayed with min and max. 3-way interaction (F(4,60) = 1.870, *P* = 0.1274); CHX x BIC (F(1,60) = 23.14, *P* < 0.0001); KCL x BIC (F(4,60) = 12.25, *P* < 0.00001); KCL x CHX (F(4,60) = 3.260, *P* = 0.0174). There were main effects of KCL and CHX but no interaction when data were collapsed over BIC: 2-way interaction KCL x CHX (F(4,70) = 0.7958, *P* = 0.5319); KCL (F(4,70) = 3.769, *P* = 0.0078); CHX (F(1,70) = 15.00, *P* = 0.0002). CHX elevated transcription in both DMSO and Bic conditions when data were collapsed over KCL (2-way interaction CHX x BIC (F(1,76) = 9.194, *P* = 0.0033); CHX(+) vs CHX(-) in Bic condition (*P* = 0.0003); in DMSO condition (*P* < 0.0001)) **(A)** All data for the experiment is displayed. **(B)** Data were collapsed over CHX: BIC x KCL Interaction (F(4,70) = 5.192, *P* = 0.001). The simple main effect of KCl on Bic treated samples was significant (F(4,35) = 6.060, *P* = 0.0008). 20mM and 30mM KCl 1° treatment resulted in significantly less induction of *Arc* pre-mRNA at the 2° stage than the mechanical controls [20mM (*P* = 0.0285); 30mM (*P* = 0.0007)]. There was no significant simple main effect of KCl on DMSO treated samples. **(C)** Samples were treated with the two-step paradigm under no treatment, +DRB (to inhibit transcription), or +DRB and +CHX (to inhibit transcription and translation together). N = 3 for all conditions. 3-way interaction (F(2,24) = 0.0407, *P* = 0.9602); BIC x KCL (F(1,24) = 2.226, *P* = 0.1487); DRB x KCL (F(2,24) = 0.2773, *P* = 0.7602); DRB x BIC (F(2,24) = 0.3183, *P* = 0.7304). There were no significant main effects of KCl or DRB, but there was a significant main effect of BIC (F(1,24) = 11.60, *P* = 0.0023). Bic treated samples induced more pERK than DMSO treated (*P* = 0.0006). **(D)** Samples were treated with and without FP during the two-step paradigm. N = 5 for all conditions. 3 way interaction (F(1,32) = 0.101, *P* = 0.753); FP x BIC (F(1,32) = 0.178, *P* = 0.676); KCl x FP (F(1,32) = 0.566, *P* = 0.458); KCl x BIC (F(1,32) = 4.250, *P* = 0.0475)). When collapsed over FP, KCL x BIC was no longer significant, but main effects of KCl and BIC were (Interaction (F(1,36) = 4.103, *P* = 0.0503); KCL (F(1,36) = 5.873, *P* = 0.021); BIC (F(1,36) = 16.22, *P* = 0.0003)). 5mM KCl had significantly more detectable pERK than 30mM KCl. Bic treated samples had significantly more detectable pERK than DMSO treated samples. † When outliers were removed, the two-way interaction KCL x BIC collapsed over FP remained significant (Interaction (F(1,34) = 12.12, *P* = 0.0014)). In Bic treated samples, there was a significant difference in pERK detected in the 5mM KCl treated samples compared with the 30mM KCL treated samples (*P* = 0.0051). **(E)** Samples were treated with 30’ 1° KCl, 1 hour washout, and 15’ 2° variable PMA to induce the MAPK pathway. N = 8 for DMSO condition, N = 4 for all PMA conditions. 2-way interaction (F(6,48) = 1.242, *P* = 0.302); main effect of KCL (F(2,48) = 1.505, *P* = 0.232); main effect of PMA (F(3, 48) = 125.2, *P* < 0.0001). All three PMA concentrations induced *Arc* pre-mRNA compared to DMSO: 1µM PMA (*P* < 0.0001), 0.1 µM PMA (*P* < 0.0001), and 0.01 µM PMA (*P* = 0.0005). Both 1µM PMA (p < 0.0001) and 0.1µM PMA (*P* < 0.0001) induced significantly more *Arc* pre-mRNA than 0.01 µM PMA. Induction of *Arc* by PMA was not depressed by 1° KCl treatment. **(F)** Samples undergoing the two-step paradigm were treated with FK from the beginning of 1° KCL (FK primary), the beginning of 2° Bic (FK secondary), or with DMSO (None). N= 4 for all cells except 5mM + Bic and 30mM FK1 + Bic, where N = 3 as values were removed due to outlying Gapdh values. For these values, the median value for the cell replaced the outlier to allow PRISM to perform the analysis. 3-way interaction (F(2,36) =1.777, *P* = 0.1837); FK x BIC (F(2,36) = 0.2023, *P* = 0.8178); FX x KCL (F(2,36) = 0.9423, *P* = 0.3991); KCL x BIC (F(1,36) = 366.5, *P* < 0.0001). Data was collapsed over FK treatment condition: Two-way interaction (F(1,42) = 326.3, *P* < 0.0001). 30mM KCl + Bic treated samples induced significantly less *Arc* pre-mRNA than 5mM KCl + Bic treated samples (*P* < 0.0001). There was no difference in induction between 30mM KCl + DMSO and 5mM KCl + DMSO samples (*P* = 0.574). **(G)** Treatment diagram for CHX, DRB and DRB/CHX, and FP experiments. **(H)** Treatment diagram for PMA experiment. **(I)** Treatment diagram for FK experiment. * indicates *P*-value < 0.05, ** < 0.01. *** < 0.001, ns = not significant.

To determine the impact of CHX, we collapsed data first over levels of KCl. CHX (+) treated samples induced significantly more transcription than CHX (-) treated samples for both Bic and DMSO treated conditions. When we collapsed data over BIC, there was no longer a significant two-way interaction between CHX x KCL. The main effect of KCL was significant, as was the main effect of CHX. Therefore, CHX elevated overall transcription (46), but was not impacted by KCl treatment. When we collapsed data over CHX, there was a significant two-way interaction between KCL and BIC (Fig 5B). As before, Bic-treated samples treated with 20mM and 30mM KCl induced significantly less *Arc* pre-mRNA than mechanical samples. There was no significant effect of KCL on DMSO-treated samples. We concluded inhibiting translation with CHX elevated overall transcription levels but did not alter the interaction between KCL and BIC. 30mM and 20mM 1° KCl pre-treatment still depressed *Arc* pre-mRNA induction by 2° Bic stimulation, and had no effect on 2° DMSO controls.

A cellular memory of mild depolarization might alternatively take the form of changes in chromatin structure or transcriptional regulation in the nucleus (2). Enhancer histone acetylation (17, 47–49) and RNA Pol pausing (29, 50) modulate the dynamics of neuronal-activity inducible genes, and could respond to previous experience. However, histone deacetylases, which negatively regulate transcription, are upregulated five or more hours after LTP induction (48). These effects are far too late to explain the short-term depressive effects described in our findings. On the other hand, RNA Pol occupancy at rapid IEGs promoters is ‘poised’ to initiate rapid transcription (29). Impaired release or depletion of paused RNA Pol at these promoters after IEG induction by our 1° KCl treatment could have depressed subsequent transcription. We therefore measured pERK induction under transcription or transcription and translation inhibition. Samples were treated with the two-step paradigm under no treatment, +DRB (to inhibit transcription), or +DRB and +CXH (to inhibit transcription and translation together). Data were analyzed with a three-way ANOVA with factors KCL (5mM or 30mM), DRB (None, +DRB, or +DRB and +CHX), and BIC (DMSO or BIC). The only significant effect was the main effect of BIC–Bic treated samples had significantly more pERK than DMSO treated samples (Fig 5C). The difference between 5mM + Bic and 30mM + Bic treated samples without DRB and CHX was not detectable in this group, and no differences were detectable when DRB and CHX were applied.

To clarify the effect of transcription inhibition, we used the transcription inhibitor flavopiridol (FP+ or FP -) in a similar set of two-step experiments (Fig 5D). In a three-way ANOVA, there were no significant interactions involving FP. When we collapsed the data over FP and analyzed with a two-way ANOVA, there was a main effect of BIC and KCl. Across all 1° treatments, Bic treated samples – as expected– had significantly more detectable pERK than DMSO. Across both 2° treatment conditions, 5mM KCl had significantly more detectable pERK than 30mM KCl. When outliers were removed from the dataset, the two-way interaction KCL x BIC was significant. In Bic treated samples, but not DMSO treated, 30mM 1° KCl significantly depressed pERK levels compared to 5mM 1° KCl (Figure 5D†). The difference between 5mM KCl and 30mM KCl pre-treatment remained under FP treatment, but seemed to disappear under DRB treatment. Therefore, we required an alternative approach to determine if nuclear events were necessary for the depression of pERK.

### Depressive effects of 1° KCl treatment are upstream of MEK-ERK signaling

Signaling cascades transmit information from a calcium influx across the cell membrane to the nucleus to initiate activity-induced transcription. Two signaling pathways that induce rapid IEGs are MAPK/ERK (51, 52) and calcineurin (53–55). Inducing these pathways independent of membrane activity or inhibiting them prior to reaching the nucleus allowed us to determine if events downstream of our manipulations (including nuclear events) were necessary for the depressive effects of 30mM KCl pre-treatments. The MAPK/ERK pathway can be induced independent of the synapse by treating cells with Tetrodotoxin (TTX) and Phorbol 12-myristate 13-acetate (PMA) (31, 56). This silences propagation of action potentials while activating PKC-dependent MAPK/ERK signaling pathway, and leads to induction of IEGs (56). If nuclear events or changes in MAPK/ERK signaling downstream of PKC are the cause of IEG depression, *Arc* transcription should remain depressed when IEG transcription is activated by PMA in the 2° stage. Therefore, we applied 1µM TTX and variable concentrations of PMA at the 2° stage instead of 5µM Bic to discover if the depression in IEG transcription after 30mM 1° KCl persisted.

Data obtained from the above-mentioned experiments were analyzed with a two-way ANOVA for factors KCL (Mechanical, 5mM, and 30mM) and PMA (DMSO, 1µM PMA, 0.1 µM PMA, or 0.01 µM PMA) (Fig 5E). All three PMA concentrations successfully induced *Arc* pre-mRNA compared to DMSO. There was no significant difference in induction between 1µM PMA and 0.1µM PMA, but 1µM PMA and 0.1µM PMA induced significantly more than 0.01 µM PMA. There was no significant interaction between KCL and, nor was there a significant main effect of KCL. Induction of *Arc* pre-mRNA by PMA was not depressed by 1° KCl treatment, indicating the cellular ‘memory’ of 1° KCl treatment is not ‘stored’ downstream of PKC in the MAPK/ERK signaling pathway.

IEG transcription is also induced via calcineurin signaling (53–55). To determine if calcineurin signaling is necessary for the effect, we treated primary cortical cells undergoing the two-step paradigm with FK506 (FK) from the beginning of 1° KCl treatment (FK primary), the beginning of 2° Bic treatment (FK secondary), or not at all (Fig 5F). FK506 blocks the activation of calcineurin (57). The data was analyzed with a three-way ANOVA for KCL (5mM or 30mM), BIC (DMSO or BIC) and FK (None, FK primary, or FK secondary). There was no significant three-way interaction, nor were there significant two-way interactions involving FK. Therefore, we collapsed the data over FK treatment to analyze KCL x BIC by two-way ANOVA. For Bic treated samples, 30mM 1° KCl still induced significantly less *Arc* pre-mRNA than 5mM 1° KCl treated samples. 2° DMSO treated samples showed no differences between KCl treatments. We concluded FK506 inhibition of calcineurin signaling did not inhibit the depression of *Arc* pre-mRNA induction by 2° 5µM Bic after 1° 30mM KCl. Therefore, calcineurin is not necessary for the depressive effect of KCl 1° treatment.

Taken together, induction of the MAPK/ERK signaling pathway independent of the synapse did not replicate the depression of *Arc* induction by 30mM KCl, nor was this depression prevented by inhibition of the calcineurin signaling pathway. Signaling and nuclear events downstream of PKC and calcineurin are not sufficient or necessary (respectively) for the effect of 1° KCl treatment on a secondary stimulus an hour later. The disappearance of the effect under DRB treatment is likely artifactual, suggesting new transcription is not necessary for the depressive effect of 1° KCl on pERK in the 2° stage, as shown by the FP data.

### 1° KCl treatment acutely silences activity and depresses subsequent spontaneous and evoked bursting

Because the depressive effects of <u>1°</u> KCl were not nuclear in nature and were upstream of PKC signaling, we next investigated whether they were related to broad measures of neuronal electrical activity. For this, we employed neuronal cultures grown on microelectrode arrays (MEAs). When the two-step stimulus paradigm was implemented on MEAs, we observed several interesting effects, summarized in Figure 6. Two-minute recordings were taken at the end of each treatment period (Fig 6A). Recordings were summarized as spike time stamps which were then analyzed for various firing related parameters, including: average spikes, average bursts, and average spikes in bursts (Fig 6B-D; averages taken from values for all electrodes across the recording period). Compared to media handing control, primary treatment with 5mM KCl enhanced the number of spikes per burst. In contrast, primary 30mM KCl treatment completely silences detectable electrical activity for the duration of measurement. By the end of the hour-long recovery period after KCl washout, some electrical activity returns, but this remained suppressed by all measured parameters. Interestingly, the secondary Bic stimulus is still then able to elicit the expected synchronous bursting response seen in control arrays, but with mildly, though statistically insignificant, attenuated spiking. Example traces from each treatment group and recording timepoint are provided (Fig 6E).

**Figure 6.**
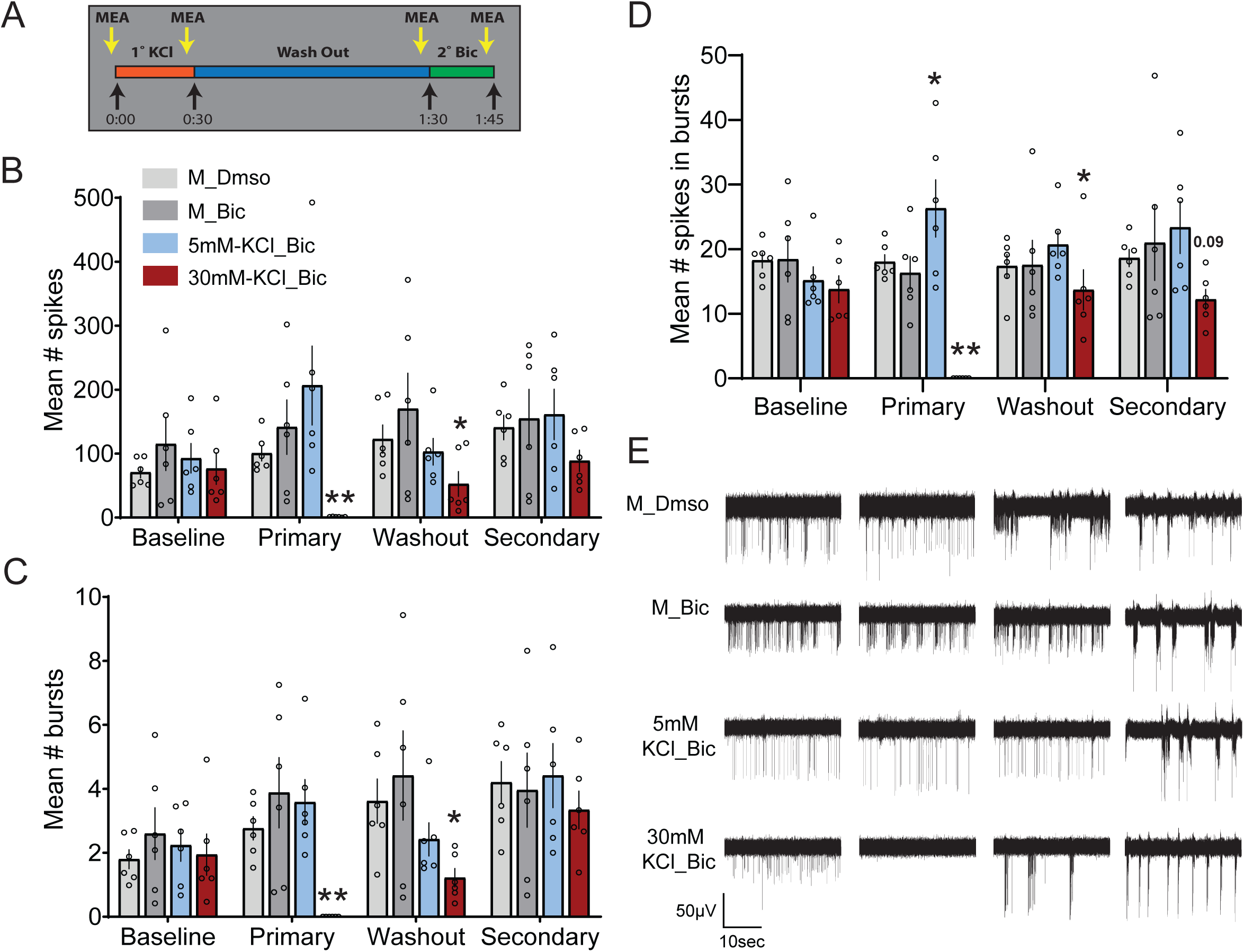
Primary treatment with 30mM KCl depresses neuronal firing and response to a secondary bicuculline stimulus. The two-step KCl treatment paradigm was implemented with neurons grown on MEAs. **(A)** Graphical summary of KCl two-step stimulus. Two-minute recordings were conducted for each treatment group at each timepoint as displayed here. **(B-D)** Recordings were summarized as spike events and analyzed for several parameters. Average number of spikes (B), average number of bursts (C), and average number of spikes in bursts (D). **(E)** Example traces of recording from each treatment group and timepoint. Note the silencing of activity observed when cells were exposed to 30mM KCl. All *P*-values were generated with a two-way ANOVA with Dunnett’s multiple comparisons test ((B) Interaction (F(9,80) = 1.312, *P* = 0.2437); Treatment (F(3,80) = 6.422, *P* = 0.0006); Timepoint (F(3,80) = 1.394, *P* = 0.2507). (C) Interaction (F(9,80) = 1.301, *P* = 0.2494); Treatment (F(3,80) = 5.580, *P* = 0.0016); Timepoint (F(3,80) = 4.324, *P* = 0.0071). (D) Interaction (F(9,80) = 2.409, *P* = 0.0179); Treatment (F(3,80) = 11.32, *P* = <0.0001); Timepoint (F(3,80) = 1.101, *P* = 0.3535.)) Comparisons were done within each timepoint, comparing each treatment group to the M_Bic array average. *N* = 6 independent culture preparations, * indicates P-value < 0.05, ** < 0.01. *** < 0.001

There are certain limitations to the initial analysis method described above. Averaging each recording across all electrodes could mask potential diverse responses at the level of individual electrodes (representing individual cells or groups of cells). This limitation may explain the sometimes-large variation in the measured firing parameters (Fig 6) and the lack of power to statistically confirm effects of 30mM KCl during the secondary stimulus. To address this, we reanalyzed the same data, but considered all electrodes individually. We initially employed this approach to investigate the effect of primary KCl treatment on spontaneous firing 1 hour after KCl washout (Fig 7). To do so, the number of spikes for each electrode was normalized to the baseline (washout-baseline, Fig 7A). We then classified electrodes as having more or less activity after primary treatment (‘up’ or ‘down’, respectively) and determined the average number of such electrodes for each replicate (Fig 7B). There were no significant differences; each treatment group consistently exhibited a similar proportion of electrodes with altered spontaneous spiking after primary KCl treatment or handling (in the case of the DMSO and M controls). Electrodes that exhibited no activity during the baseline and washout recordings were removed from the analysis (Fig 7). The magnitude of the changes in activity are represented as absolute difference, with the distribution of all electrodes, their median, and quartiles plotted in Fig 7C. Here again, we found no significant changes, though 30mM KCl treatment trended towards a greater difference in spiking from controls. However, when we considered ‘up’ or ‘down’ electrodes separately (Fig 7D-E), we observed a clear and significant attenuated effect for ‘down’ electrodes after 30mM KCl treatment (Fig 7E). Together, while there were similar proportions of electrode responses across treatment groups and no change in the magnitude of positive responses, ‘down’ electrodes –with less activity– were more suppressed when treated with 30 mM KCl.

**Figure 7.**
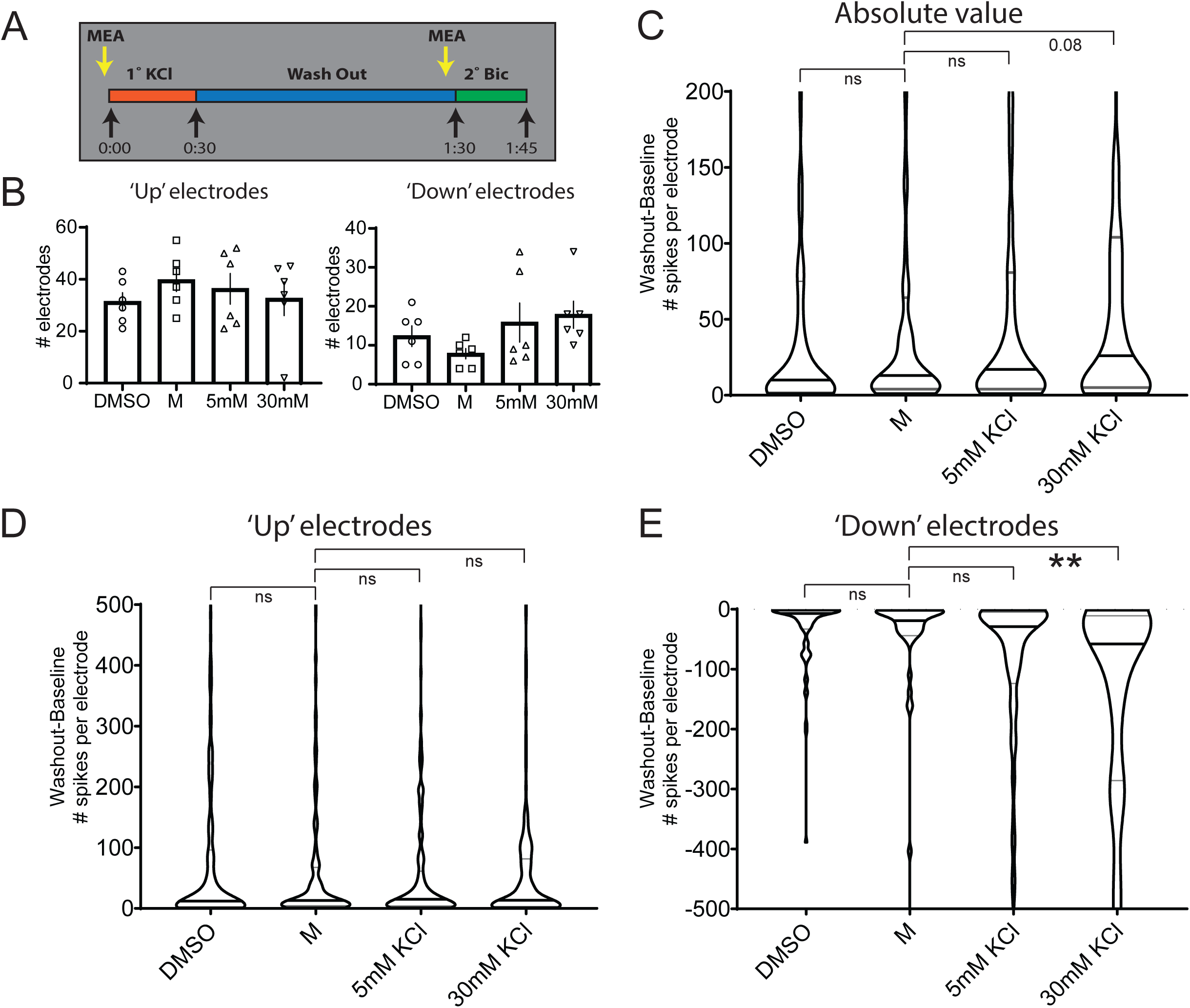
Analysis of individual electrodes from two-step KCl MEA experiments confirms a depressive effect of primary 30mM KCl treatment. Due to the observed heterogeneity of results when analyzing MEA data as averages of whole recordings (see Figure 6), we considered electrodes individually (each representing a cell or group of cells) for each treatment group across all replicates. To observe the effect of the primary treatments on spontaneous firing prior to the secondary stimulus, we compared electrodes from the washout recording to baseline (washout-baseline). **(A)** Graphical depiction of recordings used for analysis of effects on individual electrodes. The baseline electrode values were subtracted from those recorded one hour after washout of the primary treatment. **(B)** Number of electrodes classified by whether they exhibited more (up) or less (down) spiking in the washout recording compared to baseline (mean with SEM plotted). No significant changes were observed by one-way ANOVA with Dunnett’s multiple comparisons test. Electrodes up: Treatment F(3,20) = 0.5339, *P* = 0.6643. Electrodes down: Treatment F(3,20) = 1.667, *P* = 0.2060. **(C)** The absolute value of individual electrode differences found between washout and baseline are plotted, with higher values indicating a greater change in the number of spontaneous spikes per electrode. Violins represent the distribution of all individual electrodes with detectable activity across all replicates. The middle line represents the median and together with the other two lines denotes the distribution quartiles. *N* = 1161 individual electrodes. Kruskal-Wallis statistic = 15.14, *P* = 0.0017 **(D)** Electrode differences that were positive, *N* = 838 individual electrodes. Kruskal-Wallis statistic = 0.4164, *P* = 0.9368. **(E)** Differences in electrode spiking that were negative, *N* = 323 individual electrodes. Kruskal-Wallis statistic = 37.50, *P* = <0.0001. Statistical tests for C-E were non-parametric Kruskal-Wallis tests with Dunn’s multiple comparison. *N* = 6 independent culture preparations, ** indicates *P*-value < 0.01.

With electrodes now classified by effect of primary treatment on spontaneous spiking (‘up’ and ‘down’), we explored the behavior of these electrode groups during secondary Bic stimulus. Given that Bic treatment induces recurrent synchronous bursting, we further classified individual electrodes by whether they detected bursting after Bic treatment (for clarity, sample designations and filtering are detailed in Fig 8B). Figure 8A displays the average number of electrodes under each classification for each treatment group. Under control conditions, electrodes that respond to Bic with bursting tend to have also had elevated activity during the washout period. Interestingly, when cells were exposed to 30mM 1° KCl this biasing is significantly reduced and fewer electrodes exhibit bursting regardless of their washout classification. Overall, 30mM KCl treatment silences neuronal activity during its application (Fig 6B-E), suppresses subsequent spontaneous activity after washout (Fig 6B-D, 7E), and attenuates Bic-evoked activity in the form of bursting (Fig 8A).

**Figure 8.**
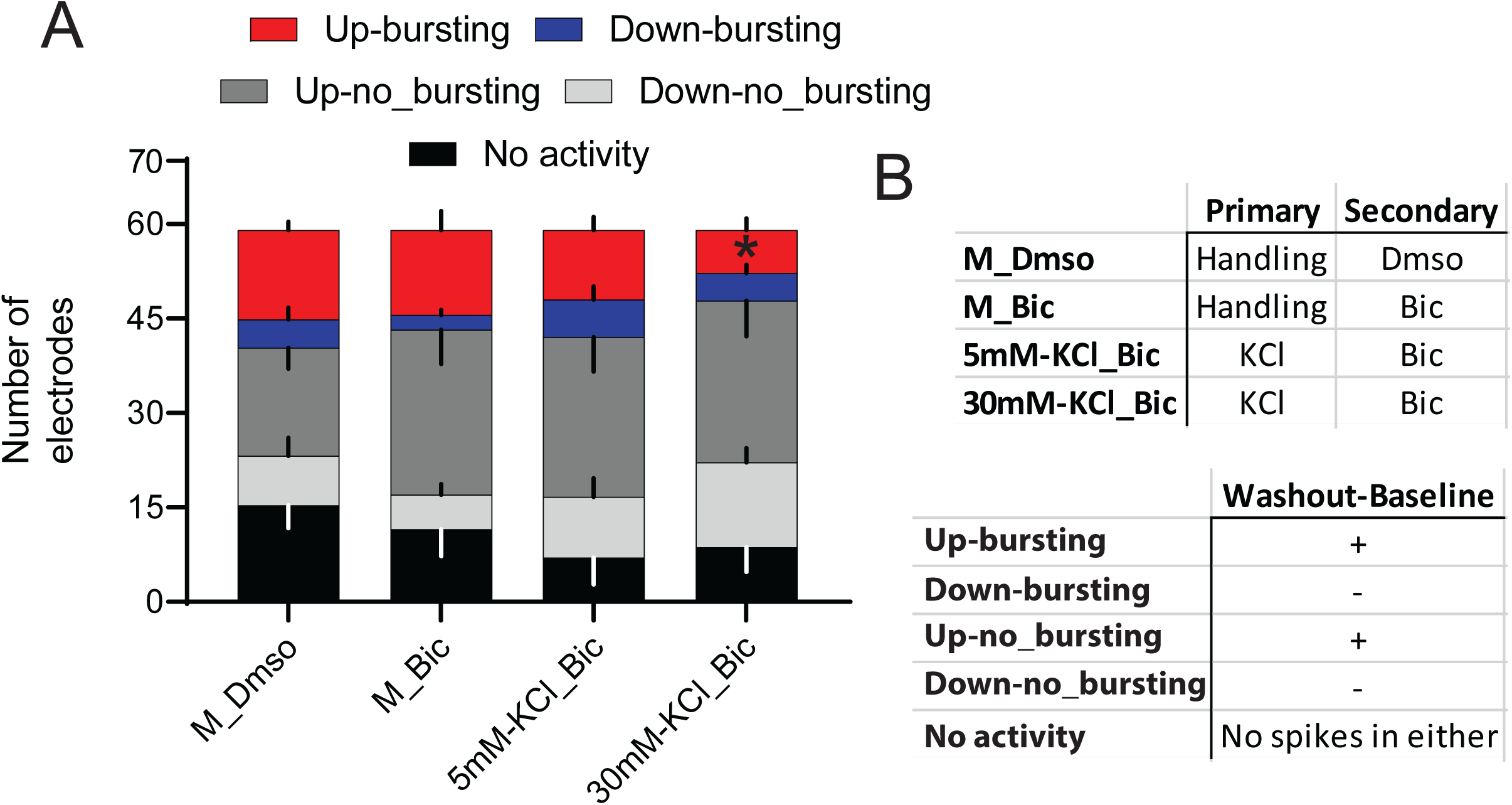
Electrodes exhibiting elevated spiking during washout are biased towards bursting during secondary bicuculline stimulus. Primary treatment with 30mM KCl reduces this bias. After classifying electrodes as exhibiting increased or decreased spiking due to primary treatment (see Figure 7), we asked what happens to these electrodes during secondary bicuculline stimulus. **(A)** The average distributions of electrode activity during secondary bicuculline treatment, broken down by status during the washout period and by whether or not the electrode subsequently displayed bursting activity at the time of secondary recording. **(B)** Tabular descriptions of labels for individual electrode data groupings referenced in (A) by treatment (upper) and washout/secondary filtering (lower). Two-way ANOVA with Dunnett’s multiple comparisons test was conducted to generate *P*-values. Interaction (F(3,40) = 2.190, *P*= 0.1042); Treatment (F(3,40) = 1.397, *P* = 0.2578); Washout difference (F(1,40) = 27.08, *P* = <0.0001). Each filtered category (Up-bursting, etc.) was compared to the corresponding M_Bic group. *N*=6 independent culture preparations. * indicates *P*-value < 0.05.

## Discussion

The activity history of a neuron impacts downstream neuronal functions (11, 23). Such history can include subthreshold graded potential changes, but effects of such mild activity remain largely unexplored. In our current study, we found mild depolarization with low concentrations of extracellular potassium chloride (KCl), 1) induced rIEG transcription and 2) depressed subsequent transcriptional, signaling (pERK), and electrical responses to synaptic stimulation an hour later. This effect occurred under 1° KCl treatments as long as 30’ and as short as 1’. When 1’ of 1° stimulus was applied, the depressive effect on transcription lasted for at least one hour, but not more than 2 hours, suggesting this mild depolarization-induced depressive effect is transient. The mechanism of this effect was not dependent on *de novo* transcription, translation, or CaN signaling, and occurred upstream of PKC in the MAPK pathway. Furthermore, MEA studies revealed neurons treated with 30mM KCl have completely attenuated spiking and bursting activity during the treatment. Activity resumed when KCl was washed out, but fewer electrodes responded to a subsequent stimulus by bursting.

Previously, we have shown that different activity patterns induce differing IEG transcription profiles (11). However, it remained unknown whether differing strengths of graded depolarization has an analogous transcriptional effect, if any at all. 50-55 mM KCl is considered to induce full strength depolarization in primary neurons (membrane potential reverses), with gene induction comparable to synaptic stimulation with 50 µM Bic. Therefore, we used low doses of 10, 20, and 30 mM KCl to investigate whether modest depolarization differentially induced rapid IEGs. Such low doses depolarize the membrane potential from the resting potential up to −45 mV (58–60). All three doses were able to induce transcription, but they did not always induce the same IEGs. Furthermore, in several cases, 20 mM KCl induced transcription while 30 mM did not (Fig. 1). Visual examination of our clustering diagram suggested linearly increasing the dose of KCl did not necessarily linearly increase IEG induction. We conclude that mild depolarization can generate robust transcription of many rapid IEGs, and the strength of such depolarization could trigger different rapid IEG transcription profile.

In addition to induction of depolarization strength-specific profiles of rapid IEGs, mild depolarization also impacted responses to a secondary synaptic stimulation. Such activity history had a depressive effect on subsequent transcription and pERK levels induced at a later time. To understand the underlying mechanism of such mild depolarization-induced ‘cellular memory’, we first investigated whether the depressive effect resulted from *de novo* transcription and/or translation. Interrupting translation with CHX–a translation inhibitor– in our two-step paradigm did not eliminate the depression of *Arc* transcription at the 2° stage. Therefore, the effects of 1° KCl after one hour is not dependent upon new protein synthesis induced by the 1° treatment. In our studies, the effects of interrupting transcription were less clear. We used pERK levels as readout for this assay upstream of transcription. DRB rescued pERK levels otherwise depressed by 30 mM 1° KCl treatment, but FP did not. While DRB and FP are both inhibitors of CDK9 within positive transcription elongation factor b (P-TEFb), which mediates Pol II elongation, FP is the more potent inhibitor and, unlike DRB, is not competitive with ATP (61–63). Furthermore, FP rapidly downregulates rapid IEGs, like FOS and GADD45B, within minutes of treatment (29, 61, 64). If the depressive effect of 1° KCl is dependent upon *de novo* transcription, it is surprising the more stringent inhibitor (FP) did not rescue pERK levels.

We took an alternative approach by determining if the cellular ‘memory’ effecting transcriptional depression existed downstream of the MAPK/ERK and CaN signaling cascades. If a cellular ‘memory’ existed at the level of gene regulation, we would have replicated the synaptic activity-dependent depressive effect on IEG transcription when we induced transcription extrasynaptically with PMA (to activate PKC) and TTX (to suppress synaptic activity) treatment. Instead, PMA induced IEG transcription to similar levels for all 1° KCl conditions, suggesting the depressive effect is upstream of transcriptional events. Furthermore, CaN inhibition had no effect on the depressive effect of 1° KCl treatment, indicating CaN signaling was not necessary for the effect. Therefore, any cellular ‘memory’ of the 1° KCl treatment likely exists upstream of PKC signaling, and not in the nucleus.

Findings discussed so far led us to investigate if 1° KCl impacted overall neuronal activity. Our MEA data revealed 30mM external KCl entirely silenced neuronal spikes and bursts during the application. This effect was curious but could be explained by refraction of voltage gated channels. After washout, neurons regained activity by the end of the recovery stage of the two-step paradigm, but total spikes, bursts, and spikes per burst remained depressed. During the 2° Bic stimulus, these properties recovered entirely, though spikes per burst trended to depression. Interestingly, after the wash out, we noticed a dichotomous response from our electrodes. Compared to baseline after 1° KCl treatment wash out, activity on some neurons increased (‘up’), while in others it decreased (‘down’). Similar duality in responses after KCl treatment have been reported previously (58). No significant difference was observed across treatment conditions in ‘up’ electrodes. However, for ‘down’ electrodes, 30 mM KCl treatment depressed spikes per electrode significantly more than other treatments.

Finally, we examined whether our ‘up’ and ‘down’ electrode groups experienced differential bursting during the 2° stimulus. In the 30mM KCl condition, fewer electrodes with elevated activity at the washout stage (‘up’) also experienced bursting in the 2° stage compared to controls. This depression in activity is not likely due to decreased cell viability by the 2° stage. First, we observed transcription depressed by 1’ of 30mM 1° KCl treatment and 1 hour wash out recovered after two hours; Second, total spikes, bursts, and spikes per burst recovered during the 2° stage; Third, KCl treatments between 25-40 mM KCl are known to have a protective effect on neuronal viability (65). In the first instance, we note that our transcriptional experiments used 1’ 1° treatment to investigate the duration of our depressive effects, but longer 1° KCl treatments may very well have longer lasting effects.

Potassium-mediated depolarization techniques are limited in that they afford only populational averages. Therefore, this method cannot investigate cell-specific responses to mild depolarization. Also, potassium-mediated depolarization protocols are not directly translatable to *in vivo* experiments. Nonetheless, we employed these protocols here as they are relatively simple and have reliably identified several signaling and transcriptional events that function in the intact brain in response to sensory stimuli (11, 17, 66, 67).

While our investigations with potassium chloride have not revealed the mechanism underlying the depressive effect of 1° mild depolarization, we have ruled out *de novo* transcription, translation, CaN signaling, and signaling events downstream of PKC. We suspect potassium treated neurons may be transiently tuning intrinsic excitability. Potential mechanisms of this transient effect include reorganization of membrane-associated structures. For instance, the axon initial segment (AIS), a specialized neuronal subcompartment localized at the beginning of the axon, is linked to modulation of intrinsic excitability (33, 36, 68). Prolonged global depolarization by KCl shifts the AIS away from the cell body in an L-type channel and CaN dependent manner, and decreases neuronal excitability (33). Many signaling pathways that respond to KCl-mediated depolarization also regulate intrinsic plasticity in response to ongoing activity (36). Also, the distribution and phosphorylation status of ion channels like the L-type channels alter the firing patterns of the cell (3, 25, 69). Interestingly, CA3 neurons in organotypic cultures modulate their intrinsic firing pattern depending on their history of ongoing subthreshold activity and kinase activity (25). Here, organotypic cultures of rat CA3 neurons experienced paired pulses applied at 1 Hz and repeated 500 times for a total of ∼8 minutes. This subthreshold conditioning did not require the cells to fire but elicited long lasting changes in the discharge dynamics of these neurons. This effect was reproduced using stimulation with intrasomatic injection of subthreshold depolarizing pulses, and separately in acute slice preparations. Importantly, the conditioning effect was blocked by adding protein kinase A (PKA) and protein kinase C (PKC) inhibitors, suggesting the changes are mediated by phosphorylation occurring over the few minutes of conditioning (25). In contrast to mechanisms of synaptic homeostatic plasticity which typically extend over hours and involve gene expression (70, 71), these experiments show changes after only a few minutes, in line with the faster timescales of intrinsic plasticity and our own experiments (72, 73).

The findings of this study may have implications in several aspects of neuronal function and dysfunction. For example, increases in extracellular K+ has been hypothesized to be etiologically relevant in epilepsy, migrainous scintillating scotoma, and other forms of cortical spreading depression (74–76). On the other hand, in normal physiology, if neurons are able to dynamically adjust their intrinsic excitability in response to their activity experience, previous mild depolarization could have an important impact in the memory engram allocation processes. Because a neuron is connected to many other neurons, synchronous firing and subsequent Hebbian potentiation between two neurons in an engram is likely restricted by the intrinsic excitability of connected downstream neurons. Neurons with high intrinsic excitability are more likely to be included in an engram; in corollary, neurons that are left out likely have lower intrinsic excitability (12, 14, 21, 22). Excitability and inclusion are in part determined by competition, wherein excitable neurons actively suppress surrounding competitors via intervening inhibitory neurons (14, 15). Enhancing CREB abundance and transcription (including CREB-dependent IEG transcription) also enhances neuronal excitability and engram allocation (12, 18, 21, 77). It is possible, based on our studies, that the recent past subthreshold activity of a given neuron also impacts competition by generating lower intrinsic excitability. In other words, neurons that have experienced mild depolarization may transiently depress responses to subsequent stimulation, thereby predisposing them to exclusion from new engrams. While this possibility remains to be tested in the brain, the idea of recent experiences dictating competitiveness in neuronal networks is nonetheless intriguing.

## Acknowledgements

We thank all Saha Lab members for their support and constructive criticism during experimentation and manuscript preparation.

## Author contributions

KDAR and RNS conceptualized the study. KDAR, RNS, RGP, and JSS conducted experiments. KDAR analyzed transcriptional and western data. RGP analyzed MEA data. KDAR (lead) and RNS wrote the manuscript with contributions from RGP. RNS supervised the entire project.

## References

1. Minatohara, K., Akiyoshi, M., and Okuno, H. (2016). Role of immediate-early genes in synaptic plasticity and neuronal ensembles underlying the memory trace. Front. Mol. Neurosci. 10.3389/fnmol.2015.00078

2. West, A. E., and Greenberg, M. E. (2011). Neuronal activity-regulated gene transcription in synapse development and cognitive function. Cold Spring Harb. Perspect. Biol. 10.1101/cshperspect.a005744

3. Flavell, S. W., and Greenberg, M. E. (2008). Signaling Mechanisms Linking Neuronal Activity to Gene Expression and Plasticity of the Nervous System. Annu. Rev. Neurosci. 10.1146/annurev.neuro.31.060407.125631

4. Wall, M. J., Collins, D. R., Chery, S. L., Allen, Z. D., Pastuzyn, E. D., George, A. J., Nikolova, V. D., Moy, S. S., Philpot, B. D., Shepherd, J. D., Müller, J., Ehlers, M. D., Mabb, A. M., and Corrêa, S. A. L. (2018). The Temporal Dynamics of Arc Expression Regulate Cognitive Flexibility. Neuron. 10.1016/j.neuron.2018.05.012

5. Gallo, F. T., Katche, C., Morici, J. F., Medina, J. H., and Weisstaub, N. V. (2018). Immediate early genes, memory and psychiatric disorders: Focus on c-Fos, Egr1 and Arc. Front. Behav. Neurosci. 10.3389/fnbeh.2018.00079

6. Pastuzyn, E. D., and Shepherd, J. D. (2017). Activity-dependent arc expression and homeostatic synaptic plasticity are altered in neurons from a mouse model of angelman syndrome. Front. Mol. Neurosci. 10.3389/fnmol.2017.00234

7. Nikolaienko, O., Patil, S., Eriksen, M. S., and Bramham, C. R. (2018). Arc protein: a flexible hub for synaptic plasticity and cognition. Semin. Cell Dev. Biol. 10.1016/j.semcdb.2017.09.006

8. Bloodgood, B. L., Sharma, N., Browne, H. A., Trepman, A. Z., and Greenberg, M. E. (2013). The activity-dependent transcription factor NPAS4 regulates domain-specific inhibition.Nature. 10.1038/nature12743

9. Sheng, H. Z., Fields, R. D., and Nelson, P. G. (1993). Specific regulation of immediate early genes by patterned neuronal activity. J. Neurosci. Res. 10.1002/jnr.490350502

10. Greenberg, M. E., Ziff, E. B., and Greene, L. A. (1986). Stimulation of neuronal acetylcholine receptors induces rapid gene transcription. Science (80-.). 10.1126/science.3749894

11. Tyssowski, K. M., DeStefino, N. R., Cho, J. H., Dunn, C. J., Poston, R. G., Carty, C. E., Jones, R. D., Chang, S. M., Romeo, P., Wurzelmann, M. K., Ward, J. M., Andermann, M. L., Saha, RN., Dudek, S. M., and Gray, J. M. (2018). Different Neuronal Activity Patterns Induce Different Gene Expression Programs. Neuron. 10.1016/j.neuron.2018.04.001

12. Yiu, A. P., Mercaldo, V., Yan, C., Richards, B., Rashid, A. J., Hsiang, H. L. L., Pressey, J., Mahadevan, V., Tran, M. M., Kushner, S. A., Woodin, M. A., Frankland, P. W., and Josselyn, S. A. (2014). Neurons Are Recruited to a Memory Trace Based on Relative Neuronal Excitability Immediately before Training. Neuron. 10.1016/j.neuron.2014.07.017

13. Guzowski, J. F., McNaughton, B. L., Barnes, C. A., and Worley, P. F. (2001). Imaging neural activity with temporal and cellular resolution using FISH. Curr. Opin. Neurobiol. 10.1016/S0959-4388(00)00252-X

14. Josselyn, S. A., and Tonegawa, S. (2020). Memory engrams: Recalling the past and imagining the future. Science (80-.). 10.1126/science.aaw4325

15. Morrison, D. J., Rashid, A. J., Yiu, A. P., Yan, C., Frankland, P. W., and Josselyn, S. A. (2016). Parvalbumin interneurons constrain the size of the lateral amygdala engram. Neurobiol. Learn. Mem. 10.1016/j.nlm.2016.07.007

16. Didier, S., Sauvé, F., Domise, M., Buée, L., Marinangeli, C., and Vingtdeux, V. (2018). AMP-activated protein kinase controls immediate early genes expression following synaptic activation through the PKA/CREB pathway. Int. J. Mol. Sci. 10.3390/ijms19123716

17. Kim, T. K., Hemberg, M., Gray, J. M., Costa, A. M., Bear, D. M., Wu, J., Harmin, D. A., Laptewicz, M., Barbara-Haley, K., Kuersten, S., Markenscoff-Papadimitriou, E., Kuhl, D., Bito, H., Worley, P. F., Kreiman, G., and Greenberg, M. E. (2010). Widespread transcription at neuronal activity-regulated enhancers. Nature. 10.1038/nature09033

18. Park, A., Jacob, A. D., Walters, B. J., Park, S., Rashid, A. J., Jung, J. H., Lau, J., Woolley, G. A., Frankland, P. W., and Josselyn, S. A. (2019). A time-dependent role for the transcription factor CREB in neuronal allocation to an engram underlying a fear memory revealed using a novel in vivo optogenetic tool to modulate CREB function. Neuropsychopharmacology. https://doi.org/10.1038/s41386-019-0588-0

19. Lisman, J., Cooper, K., Sehgal, M., and Silva, A. J. (2018). Memory formation depends on both synapse-specific modifications of synaptic strength and cell-specific increases in excitability. Nat. Neurosci. 10.1038/s41593-018-0076-6

20. Rashid, A. J., Yan, C., Mercaldo, V., Hsiang, H. L., Park, S., Cole, C. J., De Cristofaro, A., Yu, J., Ramakrishnan, C., Lee, S. Y., Deisseroth, K., Frankland, P. W., and Josselyn, S. A. (2016). Competition between engrams influences fear memory formation and recall. Science (80-.). 10.1126/science.aaf0594

21. Zhou, Y., Won, J., Karlsson, M. G., Zhou, M., Rogerson, T., Balaji, J., Neve, R., Poirazi, P., and Silva, A. J. (2009). CREB regulates excitability and the allocation of memory to subsets of neurons in the amygdala. Nat. Neurosci. 10.1038/nn.2405

22. Han, J. H., Kushner, S. A., Yiu, A. P., Cole, C. J., Matynia, A., Brown, R. A., Neve, R. L., Guzowski, J. F., Silva, A. J., and Josselyn, S. A. (2007). Neuronal competition and selection during memory formation. Science (80-.). 10.1126/science.1139438

23. Turrigiano, G., Abbott, L. F., and Marder, E. (1994). Activity-dependent changes in the intrinsic properties of cultured neurons. Science (80-.). 10.1126/science.8178157

24. Eilers, J., Augustine, G. J., and Konnerth, A. (1995). Subthreshold synaptic ca2+signalling in fine dendrites and spines of cerebellar Purkinje neurons. Nature. 10.1038/373155a0

25. Soldado-Magraner, S., Brandalise, F., Honnuraiah, S., Pfeiffer, M., Moulinier, M., Gerber, U., and Douglas, R. (2020). Conditioning by subthreshold synaptic input changes the intrinsic firing pattern of CA3 hippocampal neurons. J. Neurophysiol. 10.1152/jn.00506.2019

26. Alle, H., and Geiger, J. R. P. (2006). Combined analog and action potential coding in hippocampal mossy fibers. Science (80-.). 10.1126/science.1119055

27. Shu, Y., Hasenstaub, A., Duque, A., Yu, Y., and McCormick, D. A. (2006). Modulation of intracortical synaptic potentials by presynaptic somatic membrane potential. Nature. 10.1038/nature04720

28. Ludwar, B. C., Weiss, K. R., and Cropper, E. C. (2020). Background calcium induced by subthreshold depolarization modifies homosynaptic facilitation at a synapse in Aplysia. Sci. Rep. 10.1038/s41598-019-57362-2

29. Saha, R. N., Wissink, E. M., Bailey, E. R., Zhao, M., Fargo, D. C., Hwang, J. Y., Daigle, K. R., Fenn, J. D., Adelman, K., and Dudek, S. M. (2011). Rapid activity-induced transcription of Arc and other IEGs relies on poised RNA polymerase II. Nat. Neurosci. 10.1038/nn.2839

30. Yu, Y., Oberlaender, K., Bengtson, C. P., and Bading, H. (2017). One nuclear calcium transient induced by a single burst of action potentials represents the minimum signal strength in activity-dependent transcription in hippocampal neurons. Cell Calcium. 10.1016/j.ceca.2017.03.003

31. Poston, R. G., Murphy, L., Rejepova, A., Ghaninejad-Esfahani, M., Segales, J., Mulligan, K., and Saha, R. N. (2020). Certain ortho-hydroxylated brominated ethers are promiscuous kinase inhibitors that impair neuronal signaling and neurodevelopmental processes. J. Biol. Chem. 10.1074/jbc.RA119.011138

32. Bardy, C., Van Den Hurk, M., Eames, T., Marchand, C., Hernandez, R. V., Kellogg, M., Gorris, M., Galet, B., Palomares, V., Brown, J., Bang, A. G., Mertens, J., Böhnke, L., Boyer, L., Simon, S., and Gage, F. H. (2015). Neuronal medium that supports basic synaptic functions and activity of human neurons in vitro. Proc. Natl. Acad. Sci. U. S. A. 10.1073/pnas.1504393112

33. Evans, M. D., Sammons, R. P., Lebron, S., Dumitrescu, A. S., Watkins, T. B. K., Uebele, V. N., Renger, J. J., and Grubb, M. S. (2013). Calcineurin signaling mediates activity-dependent relocation of the Axon Initial segment. J. Neurosci. 10.1523/JNEUROSCI.0277-13.2013

34. Evans, M. D., Tufo, C., Dumitrescu, A. S., and Grubb, M. S. (2017). Myosin II activity is required for structural plasticity at the axon initial segment. Eur. J. Neurosci. 10.1111/ejn.13597

35. Azad, G. K., Ito, K., Sailaja, B. S., Biran, A., Nissim-Rafinia, M., Yamada, Y., Brown, D. T., Takizawa, T., and Meshorer, E. (2018). PARP1-dependent eviction of the linker histone H1 mediates immediate early gene expression during neuronal activation. J. Cell Biol. 10.1083/jcb.201703141

36. Grubb, M. S., and Burrone, J. (2010). Activity-dependent relocation of the axon initial segment fine-tunes neuronal excitability. Nature. 10.1038/nature09160

37. Pruunsild, P., Sepp, M., Orav, E., Koppel, I., and Timmusk, T. (2011). Identification of cis-elements and transcription factors regulating neuronal activity-dependent transcription of human BDNF gene. J. Neurosci. 10.1523/JNEUROSCI.4540-10.2011

38. Cohen, S. M., Suutari, B., He, X., Wang, Y., Sanchez, S., Tirko, N. N., Mandelberg, N. J., Mullins, C., Zhou, G., Wang, S., Kats, I., Salah, A., Tsien, R. W., and Ma, H. (2018). Calmodulin shuttling mediates cytonuclear signaling to trigger experience-dependent transcription and memory. Nat. Commun. 10.1038/s41467-018-04705-8

39. Greenberg, M. E., Greene, L. A., and Ziff, E. B. (1985). Nerve growth factor and epidermal growth factor induce rapid transient changes in proto-oncogene transcription in PC12 cells. J. Biol. Chem.

40. Dunn, C. J., Sarkar, P., Bailey, E. R., Farris, S., Zhao, M., Ward, J. M., Dudek, S. M., and Saha, R. N. (2017). Histone hypervariants H2A.Z.1 and H2A.Z.2 play independent and context-specific roles in neuronal activity-induced transcription of Arc/ Arg3.1 and other immediate early genes. eNeuro. 10.1523/ENEURO.0040-17.2017

41. Wang, D. O., Martin, K. C., and Zukin, R. S. (2010). Spatially restricting gene expression by local translation at synapses. Trends Neurosci. 10.1016/j.tins.2010.01.005

42. Jiang, C., and Schuman, E. M. (2002). Regulation and function of local protein synthesis in neuronal dendrites. Trends Biochem. Sci. 10.1016/S0968-0004(02)02190-4

43. Liu-Yesucevitz, L., Bassell, G. J., Gitler, A. D., Hart, A. C., Klann, E., Richter, J. D., Warren, S. T., and Wolozin, B. (2011). Local RNA translation at the synapse and in disease. J. Neurosci. 10.1523/JNEUROSCI.4105-11.2011

44. Li, S., Zhang, C., Takemori, H., Zhou, Y., and Xiong, Z. Q. (2009). TORC1 regulates activity-dependent CREB-target gene transcription and dendritic growth of developing cortical neurons. J. Neurosci. 10.1523/JNEUROSCI.2296-08.2009

45. España, J., Valero, J., Miñano-Molina, A. J., Masgrau, R., Martín, E., Guardia-Laguarta, C., Lleó, A., Giménez-Llort, L., Rodríguez-Alvarez, J., and Saura, C. A. (2010). β-amyloid disrupts activity-dependent gene transcription required for memory through the CREB coactivator CRTC1. J. Neurosci. 10.1523/JNEUROSCI.2154-10.2010

46. Greenberg, M. E., Hermanowski, A. L., and Ziff, E. B. (1986). Effect of protein synthesis inhibitors on growth factor activation of c-fos, c-myc, and actin gene transcription. Mol. Cell. Biol. 10.1128/mcb.6.4.1050

47. Chen, L. F., Gallegos, D. A., Hazlett, M. F., Yang, M. G., Kalmeta, B., Zhou, A. S., Grandl, J., West, A. E., Lin, Y. T., Gómez-Schiavon, M., Holtzman, L., Gersbach, C. A., and Buchler, N. E. (2019). Enhancer Histone Acetylation Modulates Transcriptional Bursting Dynamics of Neuronal Activity-Inducible Genes. Cell Rep. 10.1016/j.celrep.2019.01.032

48. Kyrke-Smith, M., and Williams, J. M. (2018). Bridging Synaptic and Epigenetic Maintenance Mechanisms of the Engram. Front. Mol. Neurosci. 10.3389/fnmol.2018.00369

49. Malik, A. N., Vierbuchen, T., Hemberg, M., Rubin, A. A., Ling, E., Couch, C. H., Stroud, H., Spiegel, I., Farh, K. K. H., Harmin, D. A., and Greenberg, M. E. (2014). Genome-wide identification and characterization of functional neuronal activity-dependent enhancers. Nat. Neurosci. 10.1038/nn.3808

50. Saha, R. N., and Dudek, S. M. (2013). Splitting Hares and Tortoises: A classification of neuronal immediate early gene transcription based on poised RNA polymerase II. Neuroscience. 10.1016/j.neuroscience.2013.04.064

51. Murphy, L. O., MacKeigan, J. P., and Blenis, J. (2004). A Network of Immediate Early Gene Products Propagates Subtle Differences in Mitogen-Activated Protein Kinase Signal Amplitude and Duration. Mol. Cell. Biol. 10.1128/mcb.24.1.144-153.2004

52. Bébien, M., Salinas, S., Becamel, C., Richard, V., Linares, L., and Hipskind, R. A. (2003). Immediate-early gene induction by the stresses anisomycin and arsenite in human osteosarcoma cells involves MAPK cascade signaling to Elk-1, CREB and SRF. Oncogene. 10.1038/sj.onc.1206334

53. Binder, A. K., Grammer, J. C., Herndon, M. K., Stanton, J. D., and Nilson, J. H. (2012). GnRH regulation of Jun and Atf3 requires calcium, calcineurin, and NFAT. Mol. Endocrinol. 10.1210/me.2012-1045

54. Lam, B. Y. H., Zhang, W., Enticknap, N., Haggis, E., Cader, M. Z., and Chawla, S. (2009). Inverse regulation of plasticity-related immediate early genes by calcineurin in hippocampal neurons. J. Biol. Chem. 10.1074/jbc.M901121200

55. Pyarajan, S., Matejovic, G., Pratt, J. C., Baksh, S., and Burakoff, S. J. (2008). Interleukin-3 (IL-3)-induced c-fos activation is modulated by Gab2-calcineurin interaction. J. Biol. Chem. 10.1074/jbc.C800087200

56. Poston, R. G., Dunn, C. J., Sarkar, P., and Saha, R. N. (2018). Persistent 6-OH-BDE-47 exposure impairs functional neuronal maturation and alters expression of neurodevelopmentally-relevant chromatin remodelers. Environ. Epigenetics. 10.1093/eep/dvx020

57. Powell, J. D., and Zheng, Y. (2006). Dissecting the mechanism of T-cell anergy with immunophilin ligands. Curr. Opin. Investig. Drugs

58. Leslie, K. R., Nelson, S. B., and Turrigiano, G. G. (2001). Postsynaptic depolarization scales quantal amplitude in cortical pyramidal neurons. J. Neurosci. 10.1523/jneurosci.21-19-j0005.2001

59. Furness, J. B. (1970). The effect of external potassium ion concentration on autonomic neuro-muscular transmission. Pflügers Arch. Eur. J. Physiol. 10.1007/BF00586580

60. Somjen, G. G. (2002). Ion regulation in the brain: Implications for pathophysiology. Neuroscientist. 10.1177/1073858402008003011

61. Keskin, H., Garriga, J., Georlette, D., and Graña, X. (2012). Complex effects of flavopiridol on the expression of primary response genes. Cell Div. 10.1186/1747-1028-7-11

62. Shao, W., and Zeitlinger, J. (2017). apoised RNA polymerase II inhibits new transcriptional initiation. Nat. Genet. 10.1038/ng.3867

63. Chao, S. H., and Price, D. H. (2001). Flavopiridol Inactivates P-TEFb and Blocks Most RNA Polymerase II Transcription in Vivo. J. Biol. Chem. 10.1074/jbc.M102306200

64. Lam, L. T., Pickeral, O. K., Peng, A. C., Rosenwald, A., Hurt, E. M., Giltnane, J. M., Averett, L. M., Zhao, H., Davis, R. E., Sathyamoorthy, M., Wahl, L. M., Harris, E. D., Mikovits, J. A., Monks, A. P., Hollingshead, M. G., Sausville, E. A., and Staudt, L. M. (2001). Genomic-scale measurement of mRNA turnover and the mechanisms of action of the anti-cancer drug flavopiridol. Genome Biol. 10.1186/gb-2001-2-10-research0041

65. Collins, F., and Lile, J. D. (1989). The role of dihydropyridine-sensitive voltage-gated calcium channels in potassium-mediated neuronal survival. Brain Res. 10.1016/0006-8993(89)90465-4

66. Chen, W. G., Chang, Q., Lin, Y., Meissner, A., West, A. E., Griffith, E. C., Jaenisch, R., and Greenberg, M. E. (2003). Derepression of BDNF Transcription Involves Calcium-Dependent Phosphorylation of MeCP2. Science (80-.). 10.1126/science.1086446

67. Rosen, L. B., and Greenberg, M. E. (1996). Stimulation of growth factor receptor signal transduction by activation of voltage-sensitive calcium channels. Proc. Natl. Acad. Sci. U. S. A. 10.1073/pnas.93.3.1113

68. Kuba, H., Oichi, Y., and Ohmori, H. (2010). Presynaptic activity regulates Na+ channel distribution at the axon initial segment. Nature. 10.1038/nature09087

69. Davis, M. J., Wu, X., Nurkiewicz, T. R., Kawasaki, J., Gui, P., Hill, M. A., and Wilson, E. (2001). Regulation of ion channels by protein tyrosin phosphorylation. Am. J. Physiol. -Hear. Circ. Physiol. 10.1152/ajpheart.2001.281.5.h1835

70. Lee, P. R., Cohen, J. E., Becker, K. G., and Fields, R. D. (2005). Gene expression in the conversion of early-phase to late-phase long-term potentiation. in Annals of the New York Academy of Sciences, 10.1196/annals.1342.023

71. Schaukowitch, K., Reese, A. L., Kim, S. K., Kilaru, G., Joo, J. Y., Kavalali, E. T., and Kim, T. K. (2017). An Intrinsic Transcriptional Program Underlying Synaptic Scaling during Activity Suppression. Cell Rep. 10.1016/j.celrep.2017.01.033

72. Aizenman, C. D., and Linden, D. J. (2000). Rapid, synaptically driven increases in the intrinsic excitability of cerebellar deep nuclear neurons. Nat. Neurosci. 10.1038/72049

73. Paz, J. T., Mahon, S., Tiret, P., Genet, S., Delord, B., and Charpier, S. (2009). Multiple forms of activity-dependent intrinsic plasticity in layer V cortical neurones in vivo. J. Physiol. 10.1113/jphysiol.2009.169334

74. de Curtis, M., Uva, L., Gnatkovsky, V., and Librizzi, L. (2018). Potassium dynamics and seizures: Why is potassium ictogenic? Epilepsy Res. 10.1016/j.eplepsyres.2018.04.005

75. Gupta, V. K. (2006). Migrainous scintillating scotoma and headache is ocular in origin: A new hypothesis. Med. Hypotheses. 10.1016/j.mehy.2005.11.010

76. Charles, A., and Brennan, K. C. (2009). Cortical spreading depression - New insights and persistent questions. Cephalalgia. 10.1111/j.1468-2982.2009.01983.x

77. Nonaka, M., Kim, R., Fukushima, H., Sasaki, K., Suzuki, K., Okamura, M., Ishii, Y., Kawashima, T., Kamijo, S., Takemoto-Kimura, S., Okuno, H., Kida, S., and Bito, H. (2014). Region-Specific Activation of CRTC1-CREB Signaling Mediates Long-Term Fear Memory. Neuron. 10.1016/j.neuron.2014.08.049

